# TelAP2 links TelAP1 to the telomere complex in *Trypanosoma brucei*

**DOI:** 10.1101/2024.05.22.595291

**Authors:** Nadine Weisert, Verena Majewski, Laura Hartleb, Katarina Luko, Liudmyla Lototska, Nils Christian Krapoth, Helle D. Ulrich, Christian J. Janzen, Falk Butter

**Author notes:** To whom correspondence should be addressed. Christian J. Janzen Falk Butter.

## Abstract

The extracellular parasite *Trypanosoma brucei* evades the immune system of the mammalian host by periodically exchanging its variant surface glycoprotein (VSG) coat. Hereby, only one VSG gene is transcribed from one of 15 subtelomeric so-called bloodstream form expression sites (BES) at any given timepoint, while all other BESs are silenced. VSG gene expression is altered by homologous recombination using a large VSG gene repertoire or by a so-called *in situ* switch, which activates a previously silent BES. Transcriptional activation, VSG switching and VSG silencing during developmental differentiation from the bloodstream form to the procyclic form present in the tsetse fly vector are tightly regulated. Due to their subtelomeric position, telomere-associated proteins are involved in the regulation of VSG expression. Three functional homologs of mammalian telomere complex proteins have been characterized thus far, and novel telomere-interacting proteins, such as telomere-associated protein 1 (TelAP1), have recently been identified. Here, we used mass spectrometry-based proteomics and interactomics approaches, telomere pull-down assays with recombinant material and immunofluorescence analysis to elucidate the interactions of 21 other putative TelAPs. We investigated the influence on VSG expression and showed that depletion of TelAPs does not ultimately lead to changes in VSG expression. Additionally, we examined the interaction patterns of four TelAPs with the *Tb*TRF/*Tb*TIF2/*Tb*RAP1 telomere complex by reciprocal affinity purification. We further propose that TelAP1 interacts with Tb927.6.4330, now called TelAP2, and that TelAP1 depends on this interaction to form a complex with the telomeric proteins *Tb*TRF, *Tb*TIF2 and *Tb*RAP1.

## Introduction

*Trypanosoma brucei* is an extracellular pathogen and the causative agent of African Sleeping sickness in humans and Nagana in cattle. During its life cycle, the parasite shuttles between the vertebrate host and the tsetse fly insect vector. To escape the immune response of the vertebrate host, *T. brucei* regularly switches its variant surface glycoprotein (VSG) coat, which consists of a dense layer of one single expressed VSG copy, to protect the parasite (1, 2). This mechanism is called antigenic variation and allows *T. brucei* to establish chronic infections by switching between 15 bloodstream form expression sites (BES). BESs are located in subtelomeric regions and are transcribed by RNA polymerase I in a polycistronic manner (3). Although transcription initiation starts at several BESs, transcription elongation throughout the entire BES is restricted to the active BES (4). The switching of the VSG gene is mediated by different mechanisms (5, 6). For example, *in situ* switching can occur via transcriptional inactivation of the promotor of the active BES and activation of a previously silent promoter. On the other hand, recombinational switches such as gene conversion and crossover events use a large repertoire of more than 2,500 internal VSG genes and pseudogenes located within VSG gene arrays on megabase chromosomes, intermediate chromosomes and minichromosomes to change the expressed VSG version (7). Specialized metacyclic expression sites (MES) are active in metacyclic stages in the salivary glands of the insect vector to prepare the parasite for transmission into the vertebrate host (3).

The monoallelic expression of only one VSG gene at any given time is tightly regulated because it is the major mechanism enabling the survival of the pathogen. BES transcription occurs at an extranucleolar compartment, the expression site body (ESB) (8). ESB is enriched in Pol-I foci necessary for transcription initiation of the BES and in the recently identified ESB-specific protein 1 (ESB1), an ES transcription activator needed for VSG expression (9). Recently, three additional nuclear bodies were found to colocalize with the ESB and associate with the active BES (10). These three nuclear splicing factor bodies (Cajal body, spliced leader (SL) array body and NUFIP splicing factor body) are important for efficient *trans*-splicing of the VSG mRNA. During mRNA maturation, an identical SL RNA is spliced to the 5’-end of each mRNA in trypanosomes, including VSG transcripts. Notably, to ensure high and efficient mRNA processing, the active BES interacts with the SL-RNA locus (11).

Furthermore, the VSG exclusion (VEX) complex is associated with the active BES in a transcription-dependent manner and is important for its interaction with the SL-RNA gene array. BES accessibility and transcription elongation are further regulated by genome architecture organization and histone modifications (12–14). While the active BES has an open chromatin structure, silent BESs are enriched in nucleosomes, leading to reduced DNA accessibility (15–17). Chromatin-associated factors such as TbISWI, disruptor of telomeric silencing 1b (*Tb*DOT1B), bromodomain factors 2 and 3 (BDF2 and BDF3), histone deacetylases and histone chaperones play a role in maintaining dense nucleosome status and the transcriptional attenuation of silent BESs (12, 18–21).

Due to the subtelomeric location of the BES, other important components influencing antigenic variation are telomeres and telomere-associated proteins (TelAPs). In mammals, telomeres are shielded from DNA repair mechanisms by the interaction of telomeric TTAGGG repeat DNA with the shelterin complex (22). Together, they form a telomeric loop structure with a 3’ TTAGGG overhang to limit nuclease access and to regulate telomerase activity (23). The three characterized *T. brucei* homologs of the mammalian shelterin complex, *Tb*TRF, *Tb*TIF2 and *Tb*RAP1, are involved in VSG expression site regulation. The homodimeric *Tb*TRF (a homolog of mammalian TRF2) directly binds telomeric DNA through its Myb domain, while *Tb*TIF2 and *Tb*RAP1 interact with *Tb*TRF (24–26). *Tb*TRF and TbTIF2 are important for telomere maintenance and integrity (27). Depletion of *Tb*TRF results in increased rates of gene conversion. Depletion of its interaction partner *Tb*TIF2 leads to the accumulation of subtelomeric double strand breaks (DSBs), which can trigger recombination events at the BES (28, 29). *Tb*RAP1 is essential for VSG silencing, and its depletion causes increased derepression, resulting in the simultaneous presence of multiple VSGs on the cell surface. Additionally, *Tb*TRF binds to noncoding telomeric repeat-containing RNA (TERRA), while the complex member TbRAP1 represses TERRA transcription, thus preventing subtelomeric DSBs caused by increased TERRA levels (30, 31). Therefore, the three homologs are important for balancing the necessity of telomere end stability and fragility to enable VSG switching events.

In a previous study, we identified potential TelAPs and further characterized two of these proteins, telomere-associated protein 1 (TelAP1) and PolIE (32, 33). TelAP1 colocalizes with *Tb*TRF in the nucleus, and its interaction with the telomere complex was confirmed by reciprocal Co-IP. The protein is involved in the regulation kinetics of developmental BES silencing during the transition from the bloodstream form (BSF) to the insect procyclic form (PCF). PolIE is a putative translesion polymerase localized at the nuclear periphery, and its depletion leads to deregulation of VSG expression and aberrant chromosome segregation. Furthermore, cells depleted of PolIE have longer 3’ overhangs, suggesting that PolIE is also involved in telomerase regulation (34).

Here, we show that PolIE, another translesion polymerase PrimPol like protein 2 (PPL2), and the two uncharacterized proteins Tb927.6.4330 (named TelAP2) and Tb927.9.3930/4000 (named TelAP3) interact with the telomere complex and suggest that TelAP2 is an important factor for the interaction of TelAP1 with the *Tb*TRF/*Tb*RAP1/*Tb*TIF2 complex.

## Materials and Methods

### Trypanosome cell line and cultivation

Monomorphic *Trypanosoma brucei* BSFs (Lister strain 427 antigenic type MITat 1.2, clone 221a) were cultured in HMI-9 medium supplemented with 10% heat-inactivated fetal calf serum at 37°C and 5% CO_2_ (35). *T. brucei* PCFs (strain 427) were cultured in modified SDM-79 medium supplemented with 10% heat-inactivated fetal calf serum at 27°C and 5% CO_2_ (36). Stable transfection by electroporation and drug selection were carried out as previously described (37). The cell population density was measured using a Beckman Coulter^TM^ Z2 particle count and size analyzer.

### Transgenic cell lines

#### MITat1.2 Cas9, 427 Cas9

For CRISPR/Cas9-based gene tagging, the transgenic Cas9 cell line was generated by transfection of the plasmid pJ1339 (gift from J. Sunter) into BSF and PCF WT cells. The plasmid was linearized with HindIII to integrate into the tubulin locus. The cell line constitutively expresses a T7 RNA polymerase, a tetracycline (tet) repressor and Cas9. *Mitat1.2 Cas9 TelAP2:PTP, Mitat1.2 Cas9 PTP:PPL2, Mitat1.2 Cas9 PTP:TelAP3, 427 Cas9 TelAP2:PTP, 427 Cas9 PTP:PPL2,* and *427 Cas9 PTP:PolIE*. A PCR-based method was used with primers containing 30 bp homologous sequences to the respective UTRs for recombination and the *PTP* open-reading frame, which were amplified from either p2678 for N-terminal tagging or p2706 for C-terminal tagging (38). The website Leishgedit.net was used for primer design. PCR amplification of the tagging constructs and sgRNA was carried out as previously described (39). Both alleles of the target genes were tagged.

#### SM PTP, 29-13 PTP

For ectopic expression of the *PTP* tag, pLEW100v5_PTP (40) was linearized with NotI and transfected into 29-13 PCF or SM cell lines (41). PTP expression was induced with 50 ng/ml tetracycline 24 h prior to the experiment.

#### 2T1 RNAi cells

The transgenic BSF 2T1 cell line (42) constitutively expressing a T7 RNA polymerase and tet repressor was used to generate RNAi cell lines. The target sequences for RNAi were amplified from genomic DNA using specific primers containing attB1 sites for insertion into the RNAi vector pGL2084 (43). The plasmid was linearized for transfection with AscI. RNAi was induced with 1 µg/ml tetracycline.

#### 2T1 TelAP2 RNAi TbTRF Ty1/-

For the *in situ* tagging of *Tb*TRF in 2T1 TelAP2 RNAi, a PCR-based method was used to delete one allele and to tag the second allele. A neomycin resistance cassette was amplified from pLF-13 using primers containing 80 bp homologous sequences to the *Tb*TRF UTRs to delete the first allele. For C-terminal tagging of the second allele, a Ty1 epitope and a puromycin resistance cassette were amplified from pMOTag2T (44) using primers containing 80 bp homologous sequences to the 3’UTR of *Tb*TRF.

All primer sequences are given in Table S1 and Table S1A in the supplemental material.

### IgG affinity purification

IgG affinity purification was performed as described previously (40). Briefly, 1-2 x 10^8^ BSF or PCF cells were harvested, washed (1500 x g, 10 min, 4°C) once with ice-cold wash solution (20 mM Tris HCl pH 7.7, 100 mM NaCl, 3 mM MgCl_2_, 1 mM EDTA) and once with ice-cold extraction buffer (150 mM sucrose, 150 mM KCl, 3 mM MgCl_2_, 20 mM HEPES-KOH, 0.1% Tween 20, 1 mM DTT, 10 µg/ml TLCK, and cOmplete^TM^ EDTA-free protease inhibitor cocktail (Roche)). Cells were lysed in 1 ml of extraction buffer by three freeze‒thaw cycles in liquid nitrogen and by sonication (2 cycles, 30 sec high-power pulse) using a Bioruptor Plus (Diagenode). After centrifugation (20,000 x g, 20 min, 4°C), the supernatant was stored at 4°C. IgG Sepharose 6 Fast Flow beads (20 µl, GE Healthcare) were equilibrated by one wash step using 1 ml of ice-cold TST buffer (50 mM Tris pH 7, 150 mM NaCl, 0.05% Tween 20) and two wash steps using 1 ml of ice-cold PA-150 buffer (150 mM KCl, 20 mM Tris-HCl pH 7.7, 3 mM MgCl_2_, 0.1% Tween 20, 0.5 mM DTT, 10 µg/ml TLCK,and cOmplete^TM^ EDTA-free protease inhibitor cocktail (Roche)) and added to the lysates. The mixture was incubated (2 h, 4°C, constant rotation). Beads were washed twice with 1 ml of ice-cold PA-150 buffer (500 x g, 5 min, 4°C). Proteins were eluted by adding 65 µl of NuPAGE LDS-Sample buffer (Thermo Fisher) supplemented with 100 mM DTT to the beads and heating for 10 min at 70°C. After centrifugation (100 x g, 1 min, RT), the supernatant was transferred to a new reaction tube using a Hamilton syringe.

### Immunoprecipitation

30 µl Protein G Sepharose 4 Fast Flow beads (GE Healthcare) were washed with 1 ml of PBS and twice with 1 ml of PBS/1% BSA. Beads were blocked in PBS/1% BSA (1 h, 4°C, constant rotation). Antibodies (monoclonal anti-Ty1 BB2 mouse antibody/monoclonal anti-TelAP1 2E6 mouse antibody) were added to the beads and incubated overnight (O/N) at 4°C with constant rotation. Beads were washed three times with 1 ml of PBS/0.1% BSA (500 × g, 1 min, 4°C) and resuspended in 100 µl of PBS/0.1% BSA. Beads were stored at 4°C until further use. Prior to usage, the beads were equilibrated by one wash step using 1 ml of ice-cold IP buffer (150 mM NaCl, 20 mM Tris HCl pH 8, 10 mM MgCl_2_, 0.5% NP-40, 10 µg/ml TLCK, and cOmplete^TM^ EDTA-free protease inhibitor cocktail (Roche)) and resuspended in 100 µl of IP buffer. A total of 2 x 10^8^ cells were harvested (1,500 x g, 10 min, 4°C) and washed once with 10 ml ice-cold TDB prior to resuspension of the pellet in 1 ml IP buffer. The cells were incubated for 20 min on ice and lysed by sonication (3 cycles, 30 sec high-power pulse) using a Bioruptor Plus (Diagenode). After centrifugation (10,000 x g, 10 min, 4°C), the supernatant was transferred to a new reaction tube, and the antibody-bead conjugates were added for incubation (overnight, 4°C, constant rotation). Beads were washed three times with 500 µl of IP buffer (500 x g, 1 min, 4°C) and resuspended in 65 µl of NuPAGE LDS-Sample buffer (Thermo Fisher) supplemented with 100 mM DTT. Proteins were eluted by heating the beads for 10 min at 70°C. After centrifugation (100 x g, 1 min, RT), the supernatant was transferred to a new reaction tube using a Hamilton syringe.

### Whole-cell lysates for mass spectrometry analysis

A total of 2 x 10^6^ cells were harvested (1500 g, 4°C, 10 min), washed with 1 ml of TDB buffer and resuspended in 60 µl of 1 x NuPAGE^TM^ LDS sample buffer (Thermo Fisher) supplemented with 100 mM DTT. The samples were boiled at 70°C for 10 min and stored at -20°C.

### Isolation of soluble VSGs

A total of 4 x 10^7^ cells were precooled on ice for 10 min prior to harvesting (1500 x g, 10 min, 4°C). After the cells were washed with 1 ml of TDB (1500 x g, 10 min, 4°C), the pellet was resuspended in 45 µl of sodium phosphate buffer supplemented with EDTA-free protease inhibitor cocktail (Roche) and incubated at 37°C for 5 min. After cooling on ice for 2 min, the cells were sedimented by centrifugation (14000 g, 5 min, 4°C), and the supernatant containing the soluble VSGs was transferred to a new reaction tube containing 15 µl of 4 x NuPAGE^TM^ LDS sample buffer (Thermo Fisher) supplemented with 400 mM DTT. The samples were boiled at 70°C for 10 min and stored at -20°C. Only cell lines without impaired cell growth were chosen for VSG analysis by mass spectrometry 72 h after induction of RNAi. For 9 cell lines that showed a growth phenotype 72 h postinduction, timepoints for harvesting were adjusted.

### Mass spectrometry and data analysis

Mass spectrometry analysis was performed in principle as described previously (45). Samples were separated on a Novex NuPAGE Bis-Tris 4–12% gradient gel (Thermo Fisher) in MES buffer (Thermo Fisher) for 8 min at 180 V. The gel was stained with Coomassie blue G250 dye (Carl Roth) prior to cutting each gel lane, mincing it and destaining with 50% ethanol in ABC buffer (50 mM ammonium bicarbonate, pH 8.0). The gel pieces were dehydrated with pure acetonitrile, reduced with 10 mM DTT (Sigma Aldrich) in ABC buffer at 56°C and alkylated with 50 mM iodoacetamide (Sigma Aldrich) in ABC buffer in the dark. The dried gel pieces were rehydrated in ABC buffer with 1 μg of trypsin per in-gel digestion at 37°C overnight. Subsequently, the digested peptides were desalted and stored on StageTips (46) for further analysis. Using a C18 reversed-phase column packed in-house with Reprosil C18 (Dr. Maisch GmbH), the peptides were separated along a 105 min gradient using an EasyLC 1000 UHPLC system. The column was enclosed in a column oven (Sonation) operated at 40°C, and peptides were sprayed onto a Q Exactive Plus mass spectrometer (Thermo), which was operated in data-dependent top 10 acquisition mode. The spray voltage was set to approximately 2.4 kilovolts. The acquired raw files were processed with MaxQuant (version 1.5.2.8) (47) using the *Trypanosoma brucei* protein database downloaded from TriTrypDB (TriTrypDB-8.1) and activated LFQ quantitation. Contaminants, reverse hits and protein groups that were identified only by site and protein groups with fewer than two peptides (one of which was unique) were removed prior to bioinformatics analysis. For enrichment, the median of the log2 LFQ intensity values of the replicates was calculated, and the p value was determined by a Welch t test between the IP and the control samples. The volcano plots were generated using the R environment.

### Western blot analysis and antibodies

Western blotting was carried out according to standard protocols. Briefly, whole cell lysates from 2 x 10^6^ cells were separated by 10% sodium dodecyl sulfate‒polyacrylamide gel electrophoresis (SDS‒PAGE) and transferred onto polyvinylidene difluoride (PVDF) membranes. The membranes were blocked in PBS/5% milk powder for 1 h at RT or O/N at 4°C. The membrane was incubated with antibodies diluted in PBS/0.1% Tween 20 solution for 1 h at RT. After each incubation with primary and secondary antibodies, three wash steps were performed using PBS/0.2% Tween 20 solution. Monoclonal rat anti-*Tb*TRF and monoclonal mouse anti-TelAP1 antibodies were used as previously described (32). Monoclonal mouse anti-PFR1,2 antibody L13D6 and mouse anti-Ty1 BB2 antibody (gift from Keith Gull, University of Oxford) were used as described (48). Primary antibodies were detected using IRdye 680LT- and 800CW-coupled antibodies diluted according to the manufacturer’s instructions with an Odyssey infrared scanner (LI-COR).

### Immunofluorescence analysis

A total of 1 x 10^6^ cells per sample were washed twice in 1 ml of vPBS (750 x g, RT, 1 min) and fixed for 1 min in PBS/4% paraformaldehyde for 20 min. Cells were subsequently resuspended in 500 µl of PBS and transferred to poly-L-lysin-coated coverslips. The cells were permeabilized in PBS/0.25% Triton-X-100 (5 min, RT) and blocked in PBS/3% BSA (30 min, RT). The blocking solution was removed, and primary antibodies diluted in PBS were added (1 h, RT, humidified chamber). After three washes with PBS (5 min, RT), secondary antibodies were applied (1 h, RT, humidified chamber, protected from light). After three washes with PBS, the coverslips were rinsed with dH_2_O and mounted with Vectashield with DAPI (Vector Laboratories). Images were captured using a Leica DMI 600B microscope and processed with Fiji software. Polyclonal antibodies against TelAP2 (1:200) and TelAP3 (1:200) were generated by immunizing a rabbit (TelAP2) and a guinea pig (TelAP3) with 500 µg of full-size recombinant HisMBP-TelAP2 and TelAP3-His proteins expressed in bacteria. Monoclonal rat anti-TbTRF and monoclonal mouse anti-TelAP1 antibodies were used as previously described (32). Primary antibodies were detected using Alexa Fluor 488- and Alexa Fluor 594-conjugated secondary antibodies (Thermo Fisher).

### Recombinant protein expression

TbTRF, TelAP2 and TelAP3 full-length coding sequences were amplified from genomic DNA and cloned and inserted into either the pCoofy4 expression vector (for N-terminal His_6_-MBP tagged TelAP2) or pET21a(+) expression vector (for C-terminal His_6_ tagged TelAP3). The TelAP2 pCoofy4 vector was transformed into BL21(DE3) pRare T1 cells, protein expression was induced, and the cells were lysed using Avestin. Purification was performed by using an MBPTrapHP column (GE Healthcare) and His-Select Ni Affinity Gel (Sigma).

The TelAP3 pET21a(+) vector was transformed into LOBSTR-BL21(DE3)-RIL E. coli cells. Protein expression was induced with 1 µM IPTG (overnight, RT, constant shaking at 180 rpm). The cells were harvested (2,000 x g, 20 min, 4°C), and the pellet was resuspended in lysis buffer (50 mM NaH_2_PO_4_ pH 8, 300 mM NaCl, 10 mM imidazole, 1% Triton X-100, 5 mM beta mercaptoethanol, 1x cOmplete^TM^ EDTA-free protease inhibitor cocktail (Roche)). The cells were sonicated (10 cycles, 30 sec high-power pulse) using a Bioruptor Plus (Diagenode) and centrifuged (10,000 x g, 10 min, 4°C). Proteins were purified using HisPur™ Ni-NTA resin (Thermo Fisher) according to the manufacturer’s instructions.

Large-scale recombinant expression of the MBP-His fusion proteins was performed in fermenters (Labfors). The fermenter was inoculated with an overnight preculture, and the culture was grown in 3 L of growth medium (82.7 g/L yeast extract, 0.8 g/L NaCl, 12.3 g/L K_2_HPO_4_, 12.3 g/L KH_2_PO_4_, 0.1 ml/L Antifoam 204, 0.6 g/L KOH, 22 g/L glucose, 0.8 g/L MgSO_4_, 30 mg/L kanamycin and 34 mg/L chloramphenicol) at 37°C and 800 rpm for 5.5 hours. The culture was induced with 1 mM IPTG and additionally grown for ca. 20 h at 24°C with 800 rpm agitation. The proteins were purified (>95%) using an FPLC system, and the correct protein size was determined by mass spectrometry.

### Telomere pull-down with recombinant proteins

Biotinylated telomeric and control DNA was prepared as previously published (49). In brief, 25 µg of a 10-mer repeat oligonucleotide was mixed in an equimolar ratio with its respective reverse complement oligonucleotide in annealing buffer (200 mM Tris-HCl, pH 8.0, 100 mM MgCl_2_, 1 M KCl). The mixture was brought to a final volume of 100 µl with water, heated at 80 °C for 5 min, and left to cool. Subsequently, the samples were supplemented with 55 µl H_2_O, 20 µl 10x T4 DNA ligase buffer (Thermo Fisher), 10 µl PEG 6000, 10 µl 100 mM ATP, 2 µl 1 M DTT and 5 µl T4 polynucleotide kinase (NEB, 10 U/µl, #M0201) and incubated at 37 °C for 2 h. Finally, 4 µl of T4 DNA ligase (Thermo Fisher, 5 WU/µl) was added, and the samples were incubated at RT overnight for ligation and polymerization. The ligation process was monitored with a 1% agarose gel. The samples were cleaned by phenol‒chloroform extraction: 1 vol. of H_2_O and 200 µl of phenol/chloroform/isoamyl alcohol (25:24:1; pH 8; Thermo Fisher) were added to the mixture, vortexed and centrifuged at 16,000 x g for 2 min. After centrifugation, the aqueous phase was transferred to a fresh tube, and the DNA was precipitated by the addition of 1 ml of pure ethanol and incubation at −20 °C for 30 min. Afterwards, the suspension was centrifuged at 16,000 x g for 45 min at 4 °C. The resulting DNA pellet was resuspended in 74 µl H_2_O, and 10 µl 10x Klenow-Fragment reaction buffer (Thermo Scientific), 10 µl 0.4 mM Biotin-dATP (Jena Bioscience) and 6 µl Klenow-Fragment exo-polymerase (Thermo Fisher, 5 U/µl) were added. Biotinylation was carried out by incubation at 37 °C overnight. The reaction mixture was cleaned by size-exclusion chromatography using MicroSpin Sephadex G-50 columns (GE Healthcare).

Biotinylated DNA and Dynabeads MyOne Streptavidin C1 (Thermo Scientific) were mixed with PBB buffer (50 mM Tris/HCl pH 7.5, 150 mM NaCl, 0.5% IGEPAL CA-630, 5 mM MgCl_2_, 1 mM DTT) and incubated at room temperature for 30 min on a rotating wheel. After three washes with PBB buffer, the DNA-coupled beads were resuspended in PBB buffer, and 10 μg of salmon sperm (Ambion) was added as a competitor for nonspecific DNA binding. The pulldowns were performed with the recombinantly expressed proteins at 4 °C on a rotating wheel for 90 min. Following incubation, the beads were washed three times with PBB buffer and resuspended in 1x LDS loading buffer (Thermo Fisher). For elution, the samples were boiled at 70 °C for 10 min and then separated on a 4-12% NuPAGE Bis-Tris gradient gel.

### Yeast two-hybrid assays

Yeast two-hybrid interaction assays were performed as described previously (50). Briefly, relevant coding sequences were cloned into bait (pGBT9) and prey (pGAD424) vectors and used in pairs to transform the yeast reporter strain PJ69-4α. After selection on synthetic complete medium without leucine and tryptophane (SC-LW), five colonies per combination were combined and suspended in ddH_2_O. The suspensions were spotted onto solid SC-LW, SC-LWH (lacking histidine), and SC-LWHA (lacking histidine and adenine) and incubated at 30°C for 48-72 h. Colony formation was imaged using an Epson scanner (Perfection V700 Photo, Software 3.81).

## Results and discussion

Influence of telomere-associated proteins on VSG expression.

Previously, we identified potential telomere-associated proteins of *T. brucei* by a quantitative mass spectrometry-based interactomics screen using two complementary biochemical approaches: DNA pulldown with oligonucleotides carrying telomeric repeats and immunoprecipitation with an antibody against the telomeric protein *Tb*TRF (32). Among these candidates, TelAP1 was identified as a stage-specific regulator of ES silencing during the differentiation of *T. brucei* (32), while another candidate, PolIE, could be characterized as a putative translesion polymerase that plays a role in VSG expression regulation and genomic integrity (33).

To characterize the remaining candidates more systematically for a potential effect on VSG expression, we depleted 21 of the 22 previously identified putative telomeric candidates (Table S1) in the bloodstream form and included PolIE in this list for further characterization. For seven of the candidates, i.e., MRBP1590 (Tb927.3.1590), PABP2 (Tb927.9.10770), PIF5 (Tb927.8.3560), PolIE (Tb927.11.5550), RPA2 (Tb927.5.1700), TelAP3 (Tb927.9.3930/Tb927.9.4000) and XRND (Tb927.10.6220), we were able to validate RNAi-mediated depletion of the protein level, as they were detected in our proteome analysis (Table S2, tabs S2B-S2T). Of the 21 generated strains, 9 showed a growth phenotype after depletion (Figure S1). This includes the already published growth deficiencies after depletion of PPL2 (Tb927.10.2520), TelAP2 (Tb927.6.4330), XRND (Tb927.10.6220) and PABP2 (Tb927.9.10770) (51–57) together with the newly observed reduced growth after p35 (Tb927.6.1190), p53 (Tb927.2.6100), p77 (Tb927.11.9920), p97 (Tb927.10.9780) and RPA2 depletion.

To analyze changes in the VSG surface coat upon mRNA depletion, VSGs were shed from the parasite cell surface before and 72 hours after RNAi induction. The VSG fraction was analyzed by high-resolution mass spectrometry, and VSG expression was quantified using MaxLFQ. For 18 of the 22 candidates, the expected VSG-2 still made up more than 99% of the quantified VSG expression. In the case of RPA2 and p97 depletion, low expression (approximately 1%) of previously silent VSGs could be observed. However, there was a notable increase in alternative VSG expression after depletion of the still uncharacterized protein TelAP2, which was comparable to the deregulation effect observed upon PolIE depletion (Figure 1A). Upregulation of BES-type VSG-6 upon TelAP2 knockdown has already been reported previously by Western blot (52). Our global VSG mass spectrometry analysis further revealed a marked upregulation of additional normally derepressed VSGs, mainly BES-associated VSGs such as VSG-8, VSG-9, VSG-15, VSG-17 and VSG-18 (Figure 1B). In particular, the expression of VSG-8 and VSG-15 was clearly upregulated upon TelAP2 depletion. A mild upregulation of these VSGs was already detected in the uninduced TelAP2 RNAi cell line, suggesting that leaky expression led to a subtle change in the VSG expression pattern. In addition to BES VSGs, we also detected VSGs from MESs, mainly VSG-397, VSG-531 and VSG-1954, and from internal genome loci such as VSG-322 and VSG-336. The proportion of MES VSGs after TelAP2 depletion was less than 1%, which contrasted with the pattern observed after PolIE or *Tb*RAP1 depletion, where most of the upregulated VSGs were expressed from MES (6 % or 3 %, respectively) (Figure 1B, Table S2).

**Figure 1:**
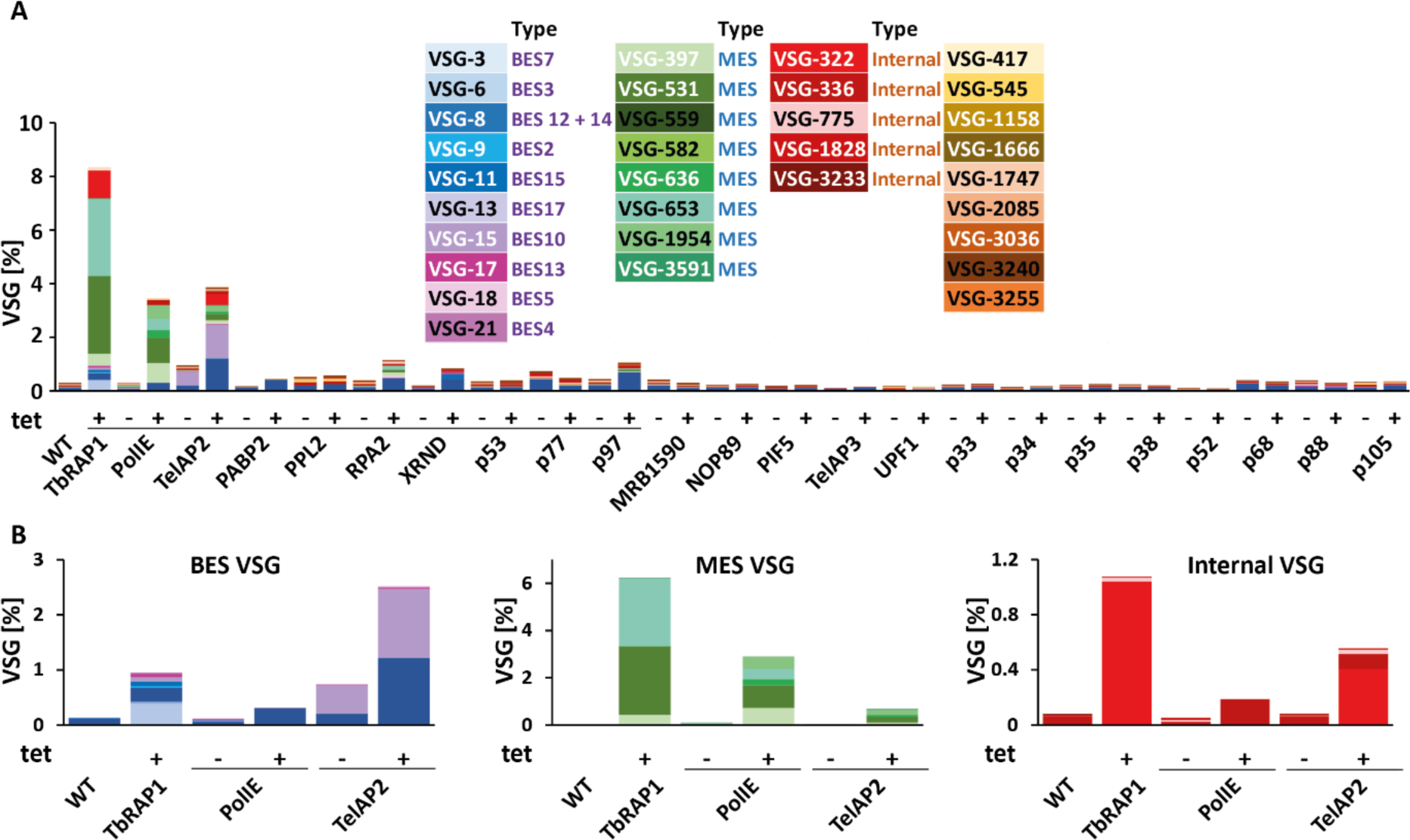
VSG expression pattern after depletion of telomere-associated protein candidates. (A) Enrichment of previously silent VSGs in induced (+) compared to uninduced (-) and wild-type (WT) cells is displayed as a percentage of the total VSG. VSG-2 (92-99% abundance) was excluded from all the graphs. RNAi was induced by adding tetracycline (tet). Cell lines that exhibited a growth phenotype after protein depletion are underlined. VSG proteins were released from the cell surface and analyzed by mass spectrometry. The published VSG expression patterns of *Tb*RAP1- and PolIE-depleted cells (33) were added for comparison. (B) Close-up of the expression patterns of previously silenced BES (left panel), MES (middle panel) and internal VSGs (right panel) of TelAP2, TbRAP1 and PolIE.

Interestingly, depletion of the previously known telomeric proteins *Tb*RAP1 and PolIE or the new candidate TelAP2 resulted in different changes in their VSG expression patterns, suggesting that different mechanisms of VSG regulation might be disrupted. In agreement with these findings, we did not observe a strong correlation between VSG deregulation and growth phenotype. For example, the severe growth phenotype after depletion of the translesion polymerase PPL2 (prim-pol like polymerase 2) had no influence on VSG expression, while depletion of PolIE deregulates VSG expression. Considering the different mechanisms of VSG switching that occur during periodical VSG exchange on the parasite surface (6), we speculate that, based on the observed VSG pattern upon knockdown, TelAP2 could be involved in BES switching rather than recombination events between BESs and MESs or internal VSG genes.

Interaction of telomere-associated proteins with the telomere complex.

Next, we investigated the composition of the telomere complex to verify that some of the putative telomeric proteins were new TelAP members. As some telomeric factors in *T. brucei* have been reported to be stage-specifically regulated (32), we generated tagged BSF and PCF strains of candidates that were previously shown to interact with TelAP1: TelAP2 (Tb927.6.4330), TelAP3 (Tb927.9.3930/Tb.927.9.4000), PPL2 (Tb927.10.2520) and PolIE (Tb927.11.5550) (32). To investigate the interactions of these four proteins, we performed affinity purification (AP) using endogenous PTP (ProtC-TEV-ProtA)-tagged proteins generated by a CRISPR-based strategy (58). To this end, prior to tagging both alleles, a BSF and a PCF cell line constitutively expressing the T7 RNA polymerase, which is necessary for *in vivo* transcription of the sgRNA, and the Cas9 enzyme, which is needed for genome editing to target the sgRNA to the gene of interest, were created (Figure S2). For TelAP2 and PPL2, it was possible to fuse the PTP tag in BSF and PCF cells, whereas for TelAP3, we could only successfully tag the gene in BSF cells and for PolIE only in PCF cells. The correct integration of the tags was confirmed by PCR analysis (Figure S3A). The growth curves of the generated cell lines confirmed that neither the constitutive expression of T7 and Cas9 nor the tagging of the proteins interfered with normal cell growth (Figure S3B). For all affinity purifications, either WT cells or cells that ectopically expressed only the PTP tag served as controls. The successful purification of the tagged proteins was confirmed by Western blotting (Figure S4). The AP experiments were analyzed by label-free quantitative mass spectrometry, and significantly enriched proteins (fold change > 2, p < 0.05, c = 0.05) are shown in the volcano plots (Figure 2). The complete TelAP2, TelAP3, PPL2 and PolIE datasets are appended (Table S3).

**Figure 2:**
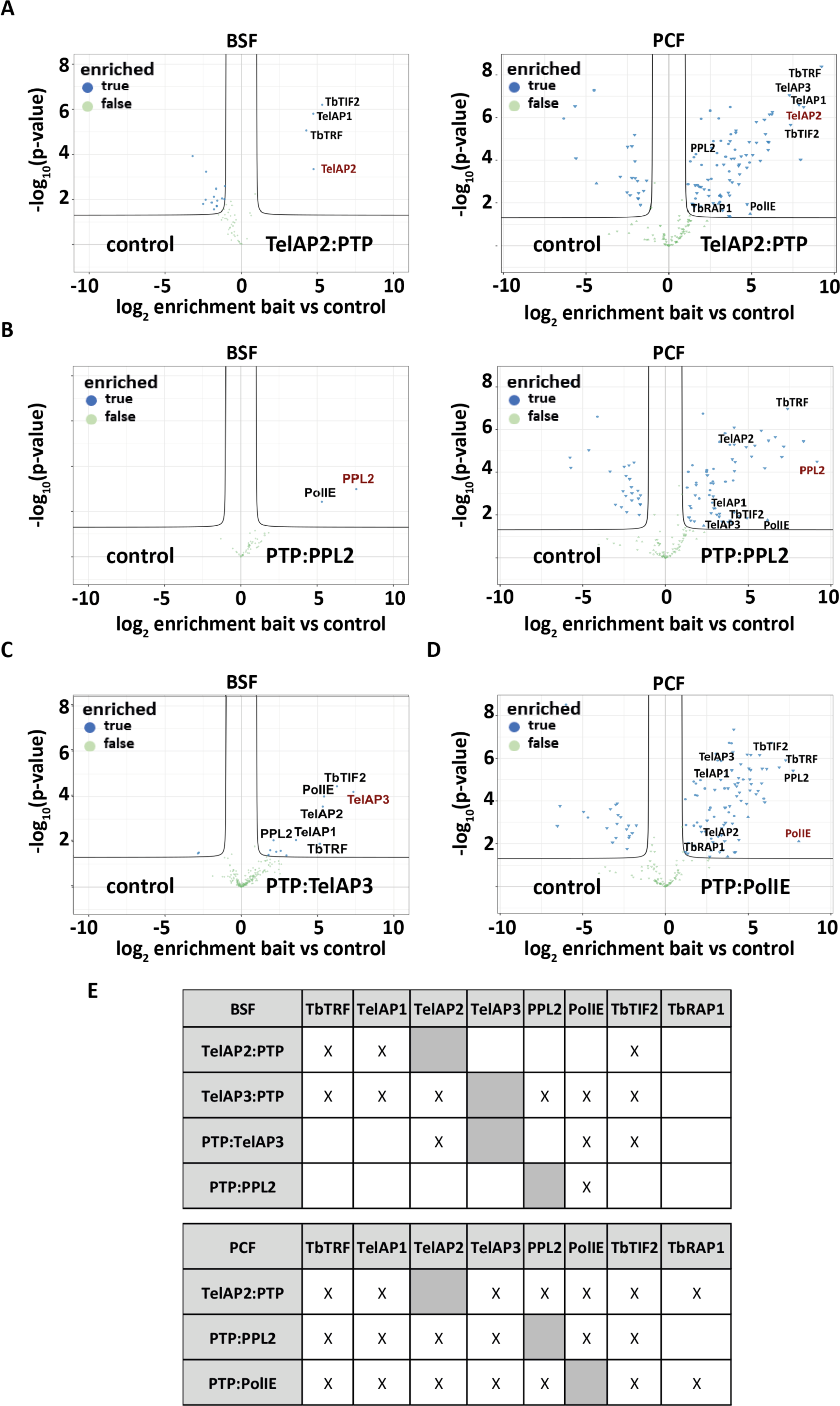
Interactome of telomeric complex proteins. (A-D) Volcano plots generated from the IP-MS data visualizing proteins copurified with the respective PTP-tagged candidates: (A) TelAP2:PTP, (B) PTP:PPL2, (C) TelAP3:PTP and (D) PTP:PolIE. The x-axis shows the log2 enrichment of quantified proteins against the control (either WT or ectopically expressed PTP), and the y-axis represents the p value (Welch t test) of replicate IPs (n=4). (E) Summary of the interactions detected in the IP experiments among selected telomeric proteins.

The number of quantified proteins ranged between 40 and 297 among the different APs. For the experiment with the lysate from BSF cells, TelAP2:PTP had only three interactors, but all of them were known telomeric proteins (*Tb*TRF, *Tb*TIF2 and TelAP1), while with lysate from the PCF strain, it interacted with 81 proteins, albeit among them also known or candidate telomeric proteins (*Tb*TRF, *Tb*TIF2, *Tb*RAP1, TelAP1, TelAP3, PPL2 and PolIE) (Figure 2A; Table S3, tab S3A, S3B). More strikingly, PTP:PPL2 only copurified with PolIE using BSF lysate, but again, 60 proteins from PCF lysate could be copurified. However, among the 60 interactors were *Tb*TRF, *Tb*TIF2, TelAP1, TelAP2, TelAP3 and PolIE (Figure 2B; Table S3, tab S3C, S3D). The AP of TelAP3:PTP, which was only taggable in BSF, revealed 11 coenriched proteins (Figure 2C), including the telomeric proteins *Tb*TRF, *Tb*TIF2, TelAP1, TelAP2, PolIE and PPL2. The AP of PTP:TelAP3 revealed 12 coenriched proteins, including *Tb*TIF2, TelAP2, TelAP3 and PolIE. Finally, the AP of PTP:PolIE, which was only available from PCF cells, contained 86 coenriched proteins, including the previously observed telomeric proteins *Tb*TRF, *Tb*TIF2, *Tb*RAP1, TelAP1, TelAP2, TelAP3 and PPL2 (Figure 2D; Table S3, tab S3G). Overall, we noted a strong difference between the number of enriched proteins depending on whether the lysate was generated from BSF or PCF. Due to these differences, we distinguished between the IPs performed with lysates from the BSF cells (TelAP2, PPL2 and TelAP3) and the PCF cells (TelAP2, PPL2 and PolIE). With respect to the BSF lysate, only 2 to 12 proteins were enriched in the APs, and most of them were nuclear. In fact, they are part of a core set of 7 telomeric proteins (*Tb*TRF, *Tb*TIF2, TelAP1, TelAP2, TelAP3, PPL2 and PolIE) that are also found in the larger interactor set of the PCF IPs (Figure 1E). Interestingly, while TbRAP1 was absent in all BSF APs, it was readily copurified in 2 (TelAP2:PTP and PTP:PolIE) of the 3 PCF AP experiments (Figure 2E). However, in the PCF AP experiments, we detected between 61 and 86 copurified proteins, a much larger set compared to BSF. Of the 85 PTP:PolIE coenriched proteins, 55 were also copurified with PTP:PPL2, while 73 were also detected in the TelAP2:PTP IP (Supplementary Table S2).

In summary, these new data further support the existence of a core set of telomeric proteins in PCF and BSF cells even beyond a previous experiment in which only TelAP2 could be coenriched with TelAP1 in PCFs (32). This is most likely due to different experimental approaches. PTP purification using a tagged cell line with a high-affinity tandem tag was described to be extremely efficient, while in the previous co-IP, a custom-made TelAP1 antibody with unknown avidity was used. Although weak and transient interactions can be missed by co-IP, we do not have any evidence for novel components of the telomere protein complex in *T. brucei* after this extensive protein‒protein interaction screen. While there were no obvious additional candidates for the core telosome, we focused on the interactions among the telosome members, especially the newly characterized proteins TelAP2 and TelAP3.

### TelAP2 is important for the interaction of TelAP1 with the telomere complex

To investigate the complex-specific interaction pattern of the two subunits TelAP2 and TelAP3, we performed immunoprecipitations with BSF cell lysates using an antibody specific for TelAP1 and depleted either TelAP2 or TelAP3. (Figure 3A and 3B; Table S4, tabs S4A, S4B). The successful immunoprecipitation of TelAP1 was confirmed by Western blotting (Figure S5A), and the samples were then subjected to quantitative mass spectrometry-based interactomics. While there were no differences upon TelAP3 knockout (Figure 3B), the depletion of TelAP2 interestingly resulted in the loss of interactions of known telomere-associated proteins, such as TbTIF2, TbTRF, TbRAP1, and PolIE, with TelAP1, suggesting that either the association of TelAP1 with the rest of the telomeric complex is dependent on TelAP2 or that reduced levels of TelAP2 destabilize TelAP1. Consistent with these findings, we detected reduced TelAP1 signal intensity upon TelAP2 depletion by immunofluorescence (Figure S6). These observations were further confirmed by reciprocal IP of TbTRF:Ty1 with lysates from TelAP2-depleted parasites. The successful IP of TbTRF was confirmed by Western blotting (Figure S5C). Despite a slightly greater enrichment of TbTRF in the control group than in the TelAP2 knockdown group, a clear reduction in TelAP2 and TelAP1 expression was observed upon depletion of TelAP2 (Figure 3C).

**Figure 3:**
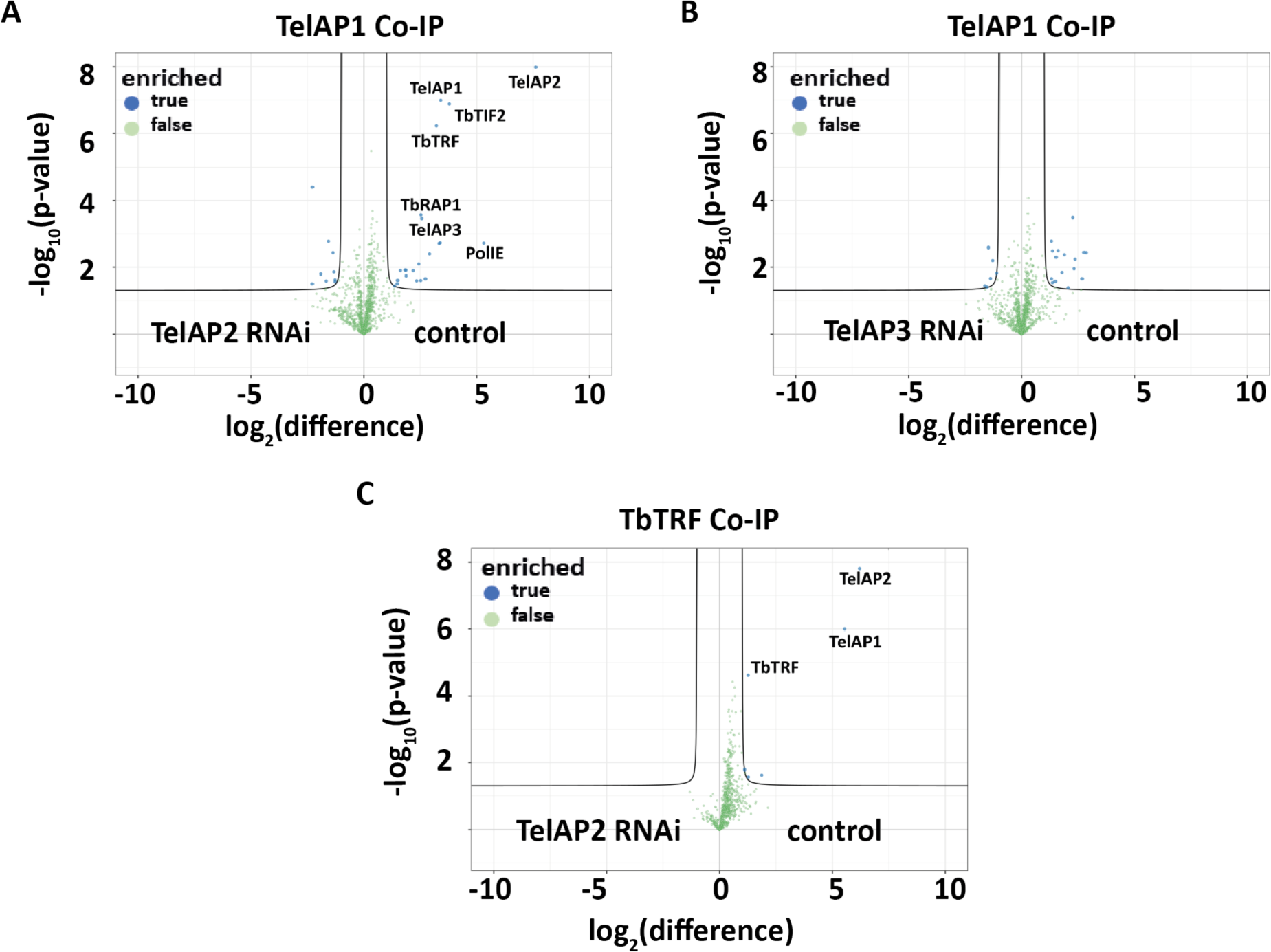
The interaction of TelAP1 with the telomere complex changes upon TelAP2 depletion. IPs were analyzed in quadruplicate by mass spectrometry using a monoclonal TelAP1 mouse antibody, and the results are displayed as a volcano plot. The x-axis shows the enrichment of quantified proteins between the uninduced cell line (control) and the cell line treated with RNAi for (A) TelAP2 or (B) TelAP3; the y-axis represents the p value (Welch t test) of replicates (n=4). **(C)** IP results of TbTRF-Ty1 in TelAP2-depleted versus control cell lysates.

To further examine whether the interaction of TelAP2 with the other subunits is DNA dependent, we performed additional immunoprecipitation experiments with TelAP2:PTP from BSF and PCF lysates with and without DNase I treatment (Figure 4A). Successful IP of the telomeric complex was confirmed by Western blotting with antibodies against the PTP-tag or TbTRF (Figure S7). Despite a very slight reduction in TelAP1 after DNase I treatment in PCFs, no other telomeric complex member showed a difference for the IP with and without DNase I treatment in BSF (Table S4, tab D) and PCF (Table S4, tab E) lysates, suggesting that TelAP2 interactions with the telomere complex do not depend on a possible interaction with DNA. In line with this, in an *in vitro* telomere pulldown assay with immobilized TTAGGG repeats versus control oligonucleotides, recombinantly expressed TelAP2 did not interact with telomeric DNA (Figure 4B). As a positive control, recombinant *Tb*TRF bound to the telomeric repeat sequence, while TelAP2 did not bind the repeat sequence directly. TelAP2 likely does not directly interact with *Tb*TRF, as it did not bind to the TTAGGG telomeric sequence when coincubated with recombinantly expressed TbTRF (Figure 4B). We thus used a yeast-2 hybrid (Y2H) assay to examine potential direct protein‒protein interactions within the telomeric complex. Two fragments each of PPL2 and PolIE and full-length TelAP1, TelAP2, TelAP3 and *Tb*TRF were inserted into bait and prey vectors and used to transform yeast cells in all possible combinations. Under highly selective conditions, only the combination of the bait TelAP1 and the prey TelAP2 allowed growth (Figure 5), thus indicating a direct interaction between the two proteins. While we cannot exclude additional interactions that might not be revealed in this assay, these data are consistent with a recruitment of TelAP1 to telomeres via TelAP2.

**Figure 4:**
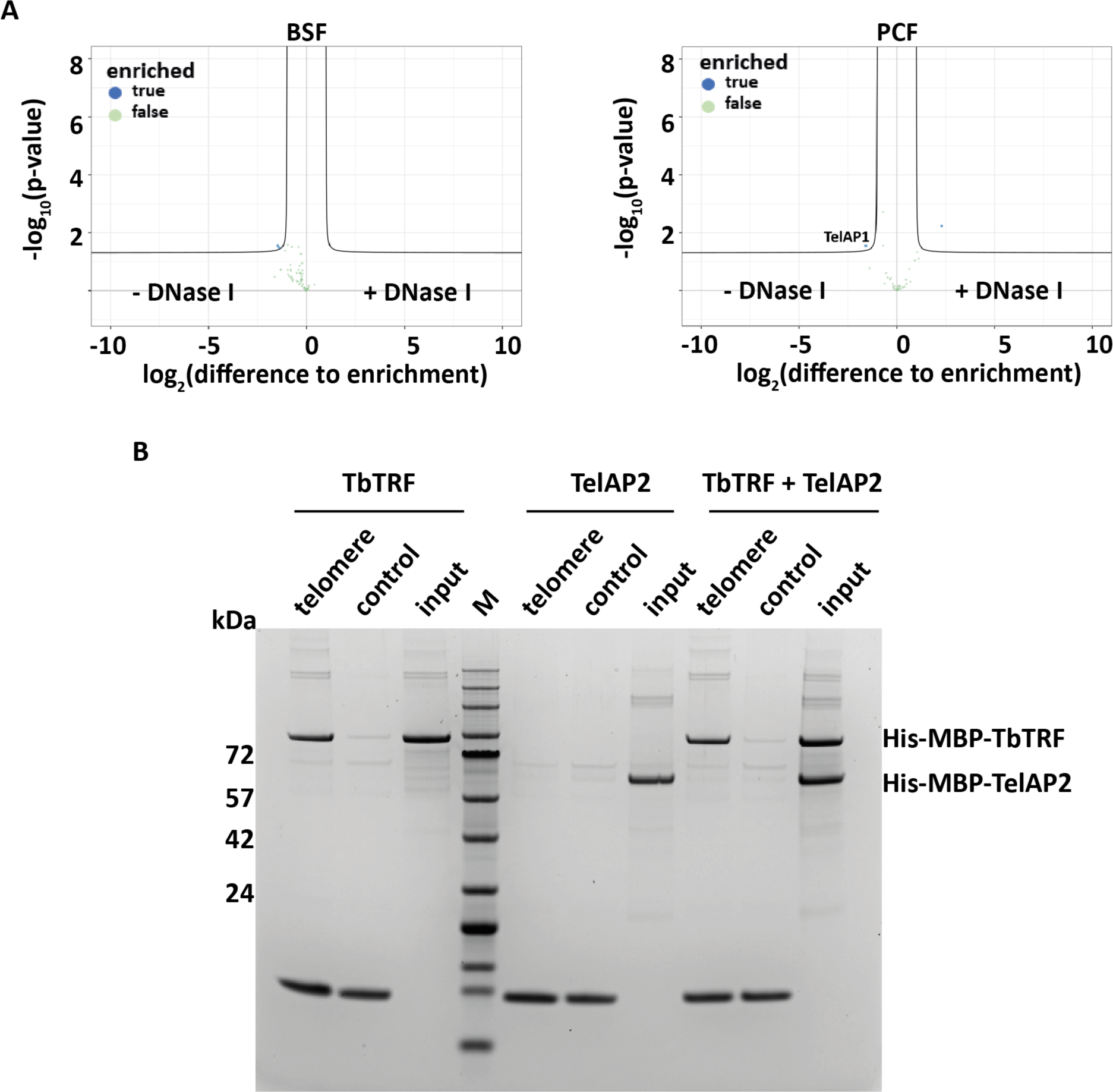
TelAP2 does not bind telomeric DNA *in vitro*. (A) Volcano plot of TelAP2 IP with and without DNase I treatment. In BSF, none of the known telomeric TelAP2 interaction partners dissociated after treatment. The x-axis shows the enrichment of proteins detected in the untreated lysates compared to the lysates treated with DNase I, and the y-axis represents the p value (Welch t test) of the quadruplicate samples. (B) *In vitro* telomere-binding assay with recombinant *Tb*TRF and TelAP2. Five micrograms of either purified His-MBP-*Tb*TRF or His-MBP-TelAP2 or a combination of *Tb*TRF and TelAP2 was used in a DNA pulldown assay. The recombinant proteins were incubated with either telomeric (TTAGGG)_n_ oligonucleotides or control (GTGATG)_n_ oligonucleotides. (M) Protein size marker.

**Figure 5:**
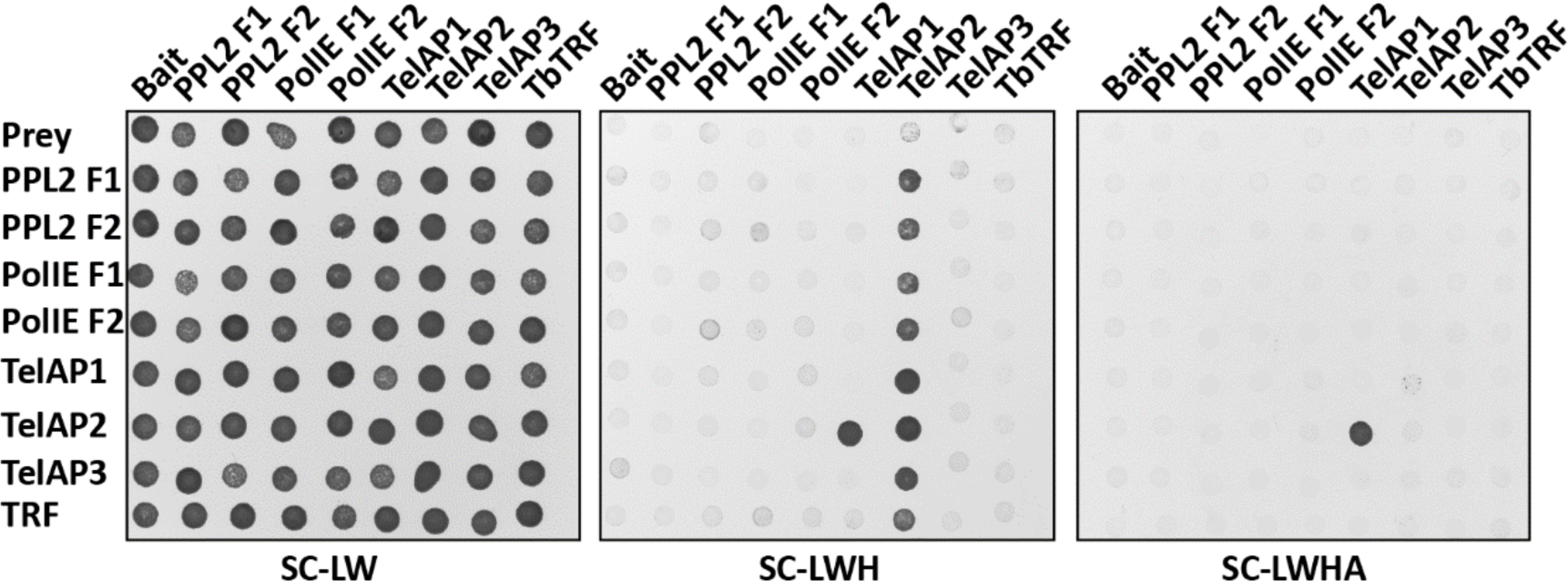
Yeast 2-hybrid assay of telomere-associated proteins shows direct interaction between TelAP1 and TelAP2. Interactions were tested in all combinations. For PPL2 and PolIE, the open reading frames were split into two fragments (F1 and F2). Yeast cells transformed with empty prey and bait vectors served as controls. Synthetic complete medium lacking leucine and tryptophane (SC-LW) ensured maintenance of the two plasmids. Growth in the absence of histdine (SC-LWH) versus histidine and adenine (SC-LWHA) indicates interactions under low versus high stringency.

## Conclusion

In this study, we used quantitative mass spectrometry to characterize previously identified potential TelAPs regarding their role in VSG expression site regulation and to identify interaction partners of these TelAPs in BSF and PCF *T. brucei* cell lines. Using an RNA interference (RNAi) screen in combination with mass spectrometry VSG analysis, we showed that one of the new candidates, TelAP2, is involved in VSG regulation. Our data suggest that *Tb*TRF, *Tb*TIF2 and *Tb*RAP1 are associated with TelAP1, TelAP2 and TelAP3. Finally, Y2H binding studies and reciprocal co-IP of TelAP1 and *Tb*TRF in TelAP2-depleted cells suggested that TelAP2 tethers TelAP1 to the rest of the telomeric protein complex via a DNA-independent interaction.

## Notes

### Competing Interest Statement

The authors have declared no competing interest.

## References

1. Cross GA. Identification, purification and properties of clone-specific glycoprotein antigens constituting the surface coat of Trypanosoma brucei. Parasitology. 1975;71(3):393–417.

2. Mugnier MR, Stebbins CE, Papavasiliou FN. Masters of Disguise: Antigenic Variation and the VSG Coat in Trypanosoma brucei. PLoS pathogens. 2016;12(9):e1005784.

3. Hertz-Fowler C, Figueiredo LM, Quail MA, Becker M, Jackson A, Bason N, et al. Telomeric expression sites are highly conserved in Trypanosoma brucei. PloS one. 2008;3(10):e3527.

4. Kassem A, Pays E, Vanhamme L. Transcription is initiated on silent variant surface glycoprotein expression sites despite monoallelic expression in Trypanosoma brucei. Proceedings of the National Academy of Sciences of the United States of America. 2014;111(24):8943–8.

5. Taylor JE, Rudenko G. Switching trypanosome coats: what’s in the wardrobe? Trends Genet. 2006;22(11):614–20.

6. Vink C, Rudenko G, Seifert HS. Microbial antigenic variation mediated by homologous DNA recombination. FEMS microbiology reviews. 2012;36(5):917–48.

7. Cross GA, Kim H-S, Wickstead B. Capturing the variant surface glycoprotein repertoire (the VSGnome) of Trypanosoma brucei Lister 427. Molecular and biochemical parasitology. 2014;195(1):59–73.

8. Navarro M, Gull K. A pol I transcriptional body associated with VSG mono-allelic expression in Trypanosoma brucei. Nature. 2001;414(6865):759–63.

9. López-Escobar L, Hänisch B, Halliday C, Ishii M, Akiyoshi B, Dean S, et al. Stage-specific transcription activator ESB1 regulates monoallelic antigen expression in Trypanosoma brucei. Nature microbiology. 2022;7(8):1280–90.

10. Budzak J, Jones R, Tschudi C, Kolev NG, Rudenko G. An assembly of nuclear bodies associates with the active VSG expression site in African trypanosomes. Nature Communications. 2022;13(1):1–18.

11. Faria J, Luzak V, Müller LS, Brink BG, Hutchinson S, Glover L, et al. Spatial integration of transcription and splicing in a dedicated compartment sustains monogenic antigen expression in African trypanosomes. Nature microbiology. 2021;6(3):289–300.

12. Alsford S, Horn D. Cell-cycle-regulated control of VSG expression site silencing by histones and histone chaperones ASF1A and CAF-1b in Trypanosoma brucei. Nucleic acids research. 2012;40(20):10150–60.

13. Pena AC, Pimentel MR, Manso H, Vaz-Drago R, Pinto-Neves D, Aresta-Branco F, et al. Trypanosoma brucei histone H1 inhibits RNA polymerase I transcription and is important for parasite fitness *in vivo*. Molecular microbiology. 2014;93(4):645–63.

14. Müller LS, Cosentino RO, Förstner KU, Guizetti J, Wedel C, Kaplan N, et al. Genome organization and DNA accessibility control antigenic variation in trypanosomes. Nature. 2018;563(7729):121–5.

15. Figueiredo LM, Cross GA. Nucleosomes are depleted at the VSG expression site transcribed by RNA polymerase I in African trypanosomes. Eukaryotic cell. 2010;9(1):148–54.

16. Narayanan MS, Rudenko G. TDP1 is an HMG chromatin protein facilitating RNA polymerase I transcription in African trypanosomes. Nucleic acids research. 2013;41(5):2981–92.

17. Aresta-Branco F, Pimenta S, Figueiredo LM. A transcription-independent epigenetic mechanism is associated with antigenic switching in Trypanosoma brucei. Nucleic Acids Res. 2016;44(7):3131–46.

18. Hughes K, Wand M, Foulston L, Young R, Harley K, Terry S, et al. A novel ISWI is involved in VSG expression site downregulation in African trypanosomes. The EMBO journal. 2007;26(9):2400–10.

19. Figueiredo LM, Janzen CJ, Cross GA. A histone methyltransferase modulates antigenic variation in African trypanosomes. PLoS biology. 2008;6(7):e161.

20. Schulz D, Mugnier MR, Paulsen EM, Kim HS, Chung CW, Tough DF, et al. Bromodomain Proteins Contribute to Maintenance of Bloodstream Form Stage Identity in the African Trypanosome. PLoS Biol. 2015;13(12):e1002316.

21. Wang QP, Kawahara T, Horn D. Histone deacetylases play distinct roles in telomeric VSG expression site silencing in African trypanosomes. Molecular microbiology. 2010;77(5):1237–45.

22. De Lange T. Shelterin: the protein complex that shapes and safeguards human telomeres. Genes & development. 2005;19(18):2100–10.

23. De Lange T. T-loops and the origin of telomeres. Nature reviews Molecular cell biology. 2004;5(4):323–9.

24. Li B, Espinal A, Cross GA. Trypanosome telomeres are protected by a homologue of mammalian TRF2. Molecular and cellular biology. 2005;25(12):5011–21.

25. Jehi SE, Wu F, Li B. Trypanosoma brucei TIF2 suppresses VSG switching by maintaining subtelomere integrity. Cell research. 2014;24(7):870–85.

26. Yang X, Figueiredo LM, Espinal A, Okubo E, Li B. RAP1 is essential for silencing telomeric variant surface glycoprotein genes in Trypanosoma brucei. Cell. 2009;137(1):99–109.

27. Jehi SE, Nanavaty V, Li B. Trypanosoma brucei TIF2 and TRF Suppress VSG Switching Using Overlapping and Independent Mechanisms. PloS one. 2016;11(6):e0156746.

28. Glover L, Alsford S, Horn D. DNA break site at fragile subtelomeres determines probability and mechanism of antigenic variation in African trypanosomes. PLoS pathogens. 2013;9(3):e1003260.

29. Thivolle A, Mehnert A-K, Tihon E, McLaughlin E, Dujeancourt-Henry A, Glover L. DNA double strand break position leads to distinct gene expression changes and regulates VSG switching pathway choice. PLoS pathogens. 2021;17(11):e1010038.

30. Saha A, Gaurav AK, Pandya UM, Afrin M, Sandhu R, Nanavaty V, et al. Tb TRF suppresses the TERRA level and regulates the cell cycle-dependent TERRA foci number with a TERRA binding activity in its C-terminal Myb domain. Nucleic acids research. 2021;49(10):5637–53.

31. Nanavaty V, Sandhu R, Jehi SE, Pandya UM, Li B. Trypanosoma brucei RAP1 maintains telomere and subtelomere integrity by suppressing TERRA and telomeric RNA: DNA hybrids. Nucleic acids research. 2017;45(10):5785–96.

32. Reis H, Schwebs M, Dietz S, Janzen CJ, Butter F. TelAP1 links telomere complexes with developmental expression site silencing in African trypanosomes. Nucleic Acids Res. 2018;46(6):2820–33.

33. Leal AZ, Schwebs M, Briggs E, Weisert N, Reis H, Lemgruber L, et al. Genome maintenance functions of a putative Trypanosoma brucei translesion DNA polymerase include telomere association and a role in antigenic variation. Nucleic acids research. 2020;48(17):9660–80.

34. Rabbani M, Tonini ML, Afrin M, Li B. POLIE suppresses telomerase-mediated telomere G-strand extension and helps ensure proper telomere C-strand synthesis in trypanosomes. Nucleic acids research. 2022;50(4):2036–50.

35. Hirumi H, Hirumi K. Continuous cultivation of Trypanosoma brucei blood stream forms in a medium containing a low concentration of serum protein without feeder cell layers. J Parasitol. 1989;75(6):985–9.

36. Brun R, Schonenberger. Cultivation and *in vitro* cloning or procyclic culture forms of Trypanosoma brucei in a semi-defined medium. Short communication. Acta Trop. 1979;36(3):289–92.

37. Burkard G, Fragoso CM, Roditi I. Highly efficient stable transformation of bloodstream forms of Trypanosoma brucei. Molecular and biochemical parasitology. 2007;153(2):220–3.

38. Kelly S, Reed J, Kramer S, Ellis L, Webb H, Sunter J, et al. Functional genomics in Trypanosoma brucei: a collection of vectors for the expression of tagged proteins from endogenous and ectopic gene loci. Mol Biochem Parasitol. 2007;154(1):103–9.

39. Beneke T, Madden R, Makin L, Valli J, Sunter J, Gluenz E. A CRISPR Cas9 high-throughput genome editing toolkit for kinetoplastids. R Soc Open Sci. 2017;4(5):170095.

40. Eisenhuth N, Vellmer T, Rauh ET, Butter F, Janzen CJ. A DOT1B/Ribonuclease H2 Protein Complex Is Involved in R-Loop Processing, Genomic Integrity, and Antigenic Variation in Trypanosoma brucei. mBio. 2021;12(6):e0135221.

41. Wirtz E, Leal S, Ochatt C, Cross GA. A tightly regulated inducible expression system for conditional gene knock-outs and dominant-negative genetics in Trypanosoma brucei. Mol Biochem Parasitol. 1999;99(1):89–101.

42. Alsford S, Horn D. Single-locus targeting constructs for reliable regulated RNAi and transgene expression in Trypanosoma brucei. Molecular and biochemical parasitology. 2008;161(1):76–9.

43. Jones NG, Thomas EB, Brown E, Dickens NJ, Hammarton TC, Mottram JC. Regulators of Trypanosoma brucei cell cycle progression and differentiation identified using a kinome-wide RNAi screen. PLoS pathogens. 2014;10(1):e1003886.

44. Oberholzer M, Morand S, Kunz S, Seebeck T. A vector series for rapid PCR-mediated C-terminal *in situ* tagging of Trypanosoma brucei genes. Molecular and biochemical parasitology. 2006;145(1):117–20.

45. Vellmer T, Hartleb L, Fradera Sola A, Kramer S, Meyer-Natus E, Butter F, et al. A novel SNF2 ATPase complex in Trypanosoma brucei with a role in H2A.Z-mediated chromatin remodelling. PLoS Pathog. 2022;18(6):e1010514.

46. Rappsilber J, Mann M, Ishihama Y. Protocol for micro-purification, enrichment, pre-fractionation and storage of peptides for proteomics using StageTips. Nat Protoc. 2007;2(8):1896–906.

47. Cox J, Mann M. Is proteomics the new genomics? Cell. 2007;130(3):395–8.

48. Bastin P, Bagherzadeh Z, Matthews KR, Gull K. A novel epitope tag system to study protein targeting and organelle biogenesis in Trypanosoma brucei. Mol Biochem Parasitol. 1996;77(2):235–9.

49. Dietz S, Almeida MV, Nischwitz E, Schreier J, Viceconte N, Fradera-Sola A, et al. The double-stranded DNA-binding proteins TEBP-1 and TEBP-2 form a telomeric complex with POT-1. Nat Commun. 2021;12(1):2668.

50. James P, Halladay J, Craig EA. Genomic libraries and a host strain designed for highly efficient two-hybrid selection in yeast. Genetics. 1996;144(4):1425–36.

51. Rudd SG, Glover L, Jozwiakowski SK, Horn D, Doherty AJ. PPL2 translesion polymerase is essential for the completion of chromosomal DNA replication in the African trypanosome. Molecular cell. 2013;52(4):554–65.

52. Glover L, Hutchinson S, Alsford S, Horn D. VEX1 controls the allelic exclusion required for antigenic variation in trypanosomes. Proc Natl Acad Sci U S A. 2016;113(26):7225–30.

53. Li C-H, Irmer H, Gudjonsdottir-Planck D, Freese S, Salm H, Haile S, et al. Roles of a Trypanosoma brucei 5′→ 3′ exoribonuclease homolog in mRNA degradation. Rna. 2006;12(12):2171–86.

54. Kramer S, Bannerman-Chukualim B, Ellis L, Boulden EA, Kelly S, Field MC, et al. Differential localization of the two T. brucei poly (A) binding proteins to the nucleus and RNP granules suggests binding to distinct mRNA pools. PloS one. 2013;8(1):e54004.

55. Shaw PL, McAdams NM, Hast MA, Ammerman ML, Read LK, Schumacher MA. Structures of the T. brucei kRNA editing factor MRB1590 reveal unique RNA-binding pore motif contained within an ABC-ATPase fold. Nucleic acids research. 2015;43(14):7096–109.

56. Delhi P, Queiroz R, Inchaustegui D, Carrington M, Clayton C. Is there a classical nonsense-mediated decay pathway in trypanosomes? PloS one. 2011;6(9):e25112.

57. Liu B, Wang J, Yildirir G, Englund PT. TbPIF5 is a Trypanosoma brucei mitochondrial DNA helicase involved in processing of minicircle Okazaki fragments. PLoS pathogens. 2009;5(9):e1000589.

58. Schimanski B, Nguyen TN, Gunzl A. Highly efficient tandem affinity purification of trypanosome protein complexes based on a novel epitope combination. Eukaryotic cell. 2005;4(11):1942–50.

