## Supplemental Figures and Table S1 for "TelAP2 links TelAP1 to the telomere complex in *Trypanosoma brucei*"

**Supplementary Table S1: List of potential telomeric proteins.** The gene IDs represent the *Trypanosoma brucei* TREU927 sequence and the *T. brucei* Lister 427 sequence, both from the tritrypdb (1). The name is either a nomenclature based on protein size (pXX) or a provided naming in the case of already characterized proteins. The nuclear enrichment score (NES) was determined through quantitative MS analysis as previously described (2).

| Gene ID | Name | Protein Description (TriTrypDB) | Localization in PCF (Tryptag; N/C-terminal tagged) | NES |
|---|---|---|---|---|
| Tb927.2.6100; Tb427_020032100 | p53 | hypothetical protein, putative | mitochondrion/ cytoplasm (points) | NA |
| Tb927.3.1590; Tb427_030015600 | MRB1590 | mitochondrial RNA binding complex 1 subunit | cytoplasm/cytoplasm | -0.16 |
| Tb927.3.2140; Tb427_030021100 | p105 | PHD finger domain protein 4 | nucleoplasm, cytoplasm | NA |
| Tb927.3.5150; Tb427_030055000 | p33 | exonuclease | cytoplasm/cytoplasm | -1.93 |
| Tb927.5.1700; Tb427_050022800 | RPA2 | replication factor A 28 kDa subunit, putative | nucleoplasm/nucleoplasm | -1.09 |
| Tb927.5.2140; Tb427_050027800 | UPF1 | regulator of nonsense transcripts 1 | cytoplasm | -0.92 |
| Tb927.6.1190; Tb427_060015500 | p35 | Plus-3 domain/Zinc finger, C3HC4 type (RING finger), putative | endocytic, cytoplasm/cell tip(posterior), basal body, pro-basal body, cytoplasm | NA |
| Tb927.6.4330; Tb427_060048400 | TelAP2 | telomere-associated protein | nucleoplasm/nucleus | 6.07 |
| Tb927.8.3560; Tb427_080040700 | PIF5 | DNA repair and recombination helicase protein | nucleolus, cytoplasm, nucleoplasm | -2.56 |
| Tb927.9.10770; Tb427_090061600 | PABP2 | polyadenylate-binding protein 2 | cytoplasm, nucleus | -0.6 |
| Tb927.9.3930/4000; Tb427_090023800/24100 | TelAP3 | hypothetical protein, conserved | nucleus, cytoplasm/nuclear lumen | 6.58 |
| Tb927.9.5020; Tb427_090028700 | p52 | HMG-box domain containing protein, putative | cytoplasm/kinetoplast | NA |
| Tb927.9.8740; Tb427_090049100 | DRBD3 | Double RNA binding domain 3 | cytoplasm, nucleoplasm/nucleoplasm, cytoplasm | 0.54 |

| Gene ID | Name | Protein Description (TriTrypDB) | Localization in PCF (Tryptag; N/C-terminal tagged) | NES |
|---|---|---|---|---|
| Tb927.10.1900; Tb427_100022200 | p88 | DNA topoisomerase IA, putative | cytoplasm/mitochondrion, antipodal sites | -2.67 |
| Tb927.10.2200; Tb427_100026200 | p38 | hypothetical protein, putative | cytoplasm/mitochondrion, kinetoplast | -3.87 |
| Tb927.10.2520; Tb427_100029500 | PPL2 | PrimPol-like protein 2 | nucleoplasm/nucleoplasm, cytoplasm | 5.54 |
| Tb927.10.4220; Tb427_100046500 | p34 | hypothetical protein, putative | cytoplasm | -3.66 |
| Tb927.10.6220; Tb427_070054600 | XRND | 5'-3' exoribonuclease D | nucleolus/nucleolus, nucleoplasm | 3.31 |
| Tb927.10.9780; Tb427_100104600 | p97 | ATP-dependent DEAD/H RNA helicase, putative | nucleolus/nucleolus | 1.74 |
| Tb927.11.16120; Tb427_110180600 | p68 | hypothetical protein, putative | endocytic, cytoplasm/kinetoplast | NA |
| Tb927.11.5550; Tb427_110060600.1 | PolIE | DNA polymerase theta | nucleus (points)/nuclear lumen, cytoplasm | 5.74 |
| Tb927.11.6790; Tb427_110073000 | NOP89 | Nucleolar protein 89 | nucleolus/nucleolus | 2.99 |
| Tb927.11.9920; Tb427_110107600 | p77 | polyubiquitin, putative | cytoplasm, flagellar cytoplasm, nuclear lumen, flagella connector, endocytic/cytoplasm | -1.55 |

**Supplementary Table S5: Enriched proteins in PCF cells.** Proteins with a nuclear localization based on TriTrypDB GO cellular component. Telomere complex proteins and TelAPs are indicated in bold. The nuclear enrichment score (NES) is included. Proteins copurified with the respective bait (PTP:PPL2, PTP:PolIE or TelAP2:PTP) are marked with an X.

| TREU927 Gene ID | Name | Protein description | NES | PPL2 | PolIE | TelAP2 |
|---|---|---|---|---|---|---|
| **Tb927.11.9870** | **TelAP1** | **telomere-associated protein 1** | **0.96** | **X** | **X** | **X** |
| Tb927.10.3200 | U2AF35 | U2 splicing auxiliary factor, putative | 1.45 | **X** | **X** | **X** |
| Tb927.10.2890 | | enolase | 1.52 | | **X** | **X** |
| Tb927.11.1900 | | T-complex protein 1, beta subunit, putative | 1.58 | | **X** | |
| Tb927.10.13720 | RBP29 | RNA-binding protein 29, putative | 2.56 | **X** | **X** | **X** |
| **Tb927.10.2520** | **PPL2** | **PrimPol-like protein 2** | **5.54** | **X** | **X** | **X** |
| **Tb927.11.5550** | **PolIE** | **DNA polymerase theta** | **5.74** | **X** | **X** | **X** |
| **Tb927.10.12850** | ***Tb*TRF** | **ttaggg binding factor** | **6.44** | **X** | **X** | **X** |
| **Tb927.9.4000** | **TelAP3** | **hypothetical protein, conserved** | **6.58** | **X** | **X** | **X** |
| Tb927.11.16130 | NDPK | nucleoside diphosphate kinase | NA | **X** | **X** | **X** |
| Tb927.10.2290 | J3 | chaperone protein DnaJ, putative | NA | **X** | **X** | **X** |
| Tb927.11.880 | CYPA | cyclophilin a | NA | **X** | **X** | **X** |
| Tb927.11.9590 | | S-adenosylhomocysteine hydrolase, putative | NA | | **X** | **X** |
| Tb927.11.4700 | | prostaglandin f synthase | NA | | **X** | **X** |
| Tb927.2.5160 | J2 | Chaperone protein DnaJ 2 | NA | **X** | **X** | **X** |
| Tb927.9.4680 | EIF4A1 | Eukaryotic initiation factor 4A-1 | NA | **X** | **X** | **X** |
| **Tb927.3.1560** | ***Tb*TIF2** | **TRF-Interacting Factor 2** | NA | **X** | **X** | **X** |
| Tb927.9.15150 | L5 | 60S ribosomal protein L5, putative | NA | | **X** | **X** |
| Tb927.3.1790 | | pyruvate dehydrogenase E1 beta subunit, putative | NA | | **X** | **X** |
| Tb927.11.9710 | RPL10A | 60S ribosomal protein L10a, putative | NA | **X** | **X** | **X** |
| Tb927.10.6070 | UMSBP1 | universal minicircle sequence binding protein 1 | NA | **X** | **X** | **X** |
| Tb927.10.4120 | RPL30 | 60S ribosomal protein L30 | NA | | **X** | |
| Tb927.9.5320 | | nucleolar RNA binding protein, putative | NA | | **X** | |
| Tb927.10.5770 | VCP | Valosin-containing protein | NA | **X** | **X** | **X** |
| **Tb927.6.4330** | **TelAP2** | **telomere-associated protein** | **NA** | **X** | **X** | **X** |
| Tb927.10.4570 | | elongation factor 2 | NA | | **X** | **X** |
| Tb927.3.1120 | RTB2 | GTP-binding nuclear protein rtb2, putative | NA | **X** | **X** | **X** |
| Tb927.8.900 | TSR1 | splicing factor TSR1 | NA | | **X** | **X** |
| **Tb927.11.370** | ***Tb*RAP1** | **repressor activator protein 1** | **NA** | | **X** | **X** |
| Tb927.11.11370 | RACK1 | receptor for activated C kinase 1 | NA | **X** | **X** | **X** |

| TREU927 Gene ID | Name | Protein description | NES | PPL2 | PolIE | TelAP2 |
|---|---|---|---|---|---|---|
| Tb927.11.14000 | NRBD1 | nuclear RNA binding domain 1 | NA | | **X** | |
| Tb927.7.2070 | | heat shock protein DNAJ, putative | NA | | **X** | **X** |
| Tb927.4.2040 | ALBA3 | DNA/RNA-binding protein Alba 3 | NA | | **X** | **X** |
| Tb927.7.1730 | | 60S ribosomal protein L7, putative | NA | | **X** | **X** |
| Tb927.10.3210 | | delta-1-pyrroline-5-carboxylate dehydrogenase, putative | NA | | **X** | |
| Tb927.10.14550 | HEL67 | ATP-dependent RNA helicase HEL67 | NA | **X** | **X** | **X** |
| Tb927.8.3750 | NOP56 | nucleolar protein 56 | NA | | **X** | |
| Tb927.9.11410 | | 60S ribosomal protein L23, putative | NA | | **X** | **X** |
| Tb927.10.5340 | RPS18 | 40S ribosomal protein S18, putative | NA | **X** | **X** | **X** |
| Tb927.10.8020 | POP | prolyl endopeptidase | NA | | **X** | |
| Tb927.9.10770 | PABP2 | polyadenylate-binding protein 2 | NA | **X** | **X** | **X** |
| Tb927.2.340 | RHS4 | retrotransposon hot spot protein 4 (RHS4), putative | NA | | **X** | |
| Tb927.9.6070 | RPS3 | 40S ribosomal protein S3, putative | NA | | **X** | |

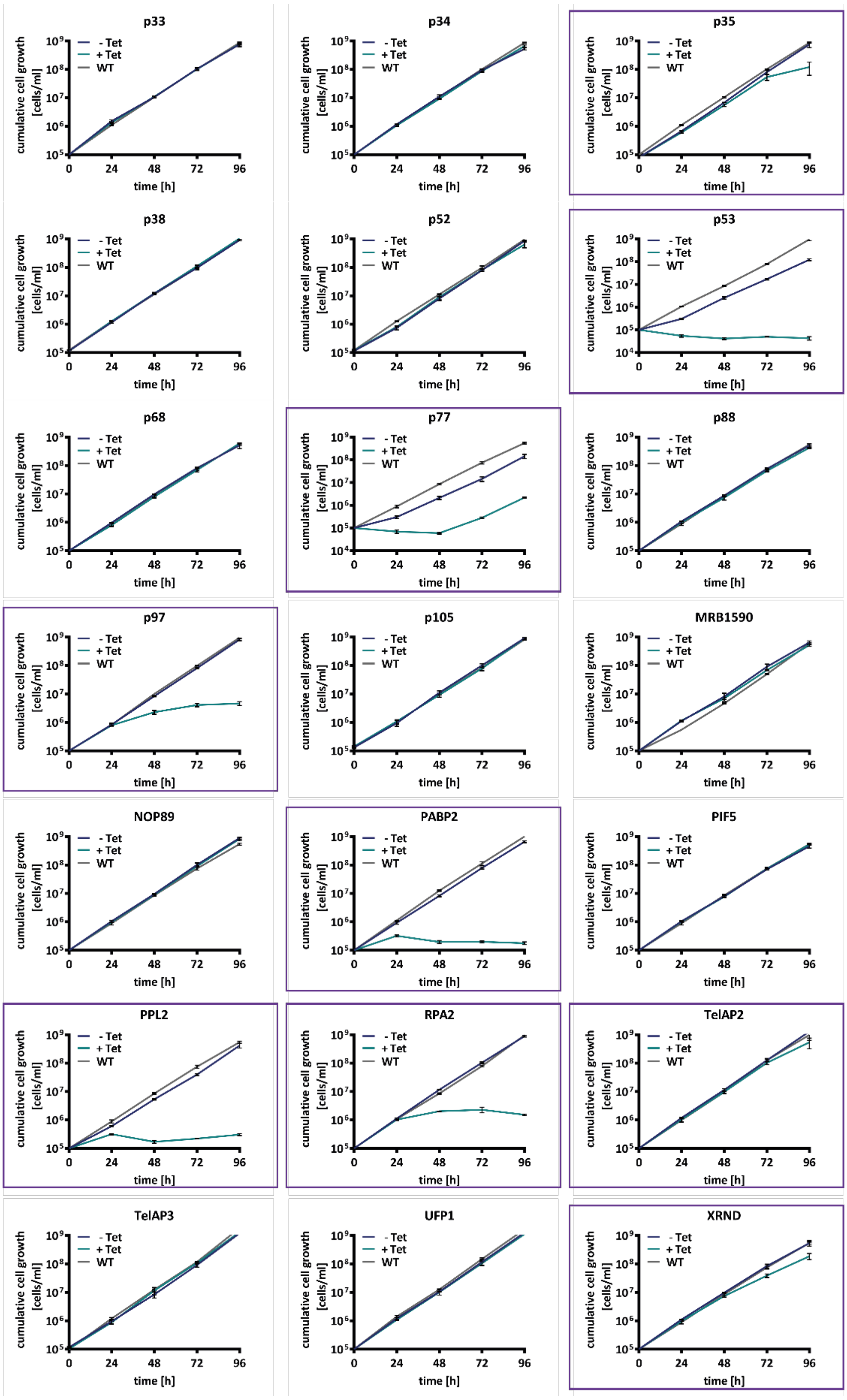


**Supplementary Figure S1: Growth curves of cell lines with depleted potential telomeric proteins via RNAi in BSF.** Measurement of the cumulative growth of wild-type cells (WT), noninduced RNAi cells (-) and cells after RNAi induction (+) using tetracycline (Tet) (n=3). Error bars denote the standard deviation (SD). Nine out of 21 cells exhibited a growth phenotype (indicated by a purple square).

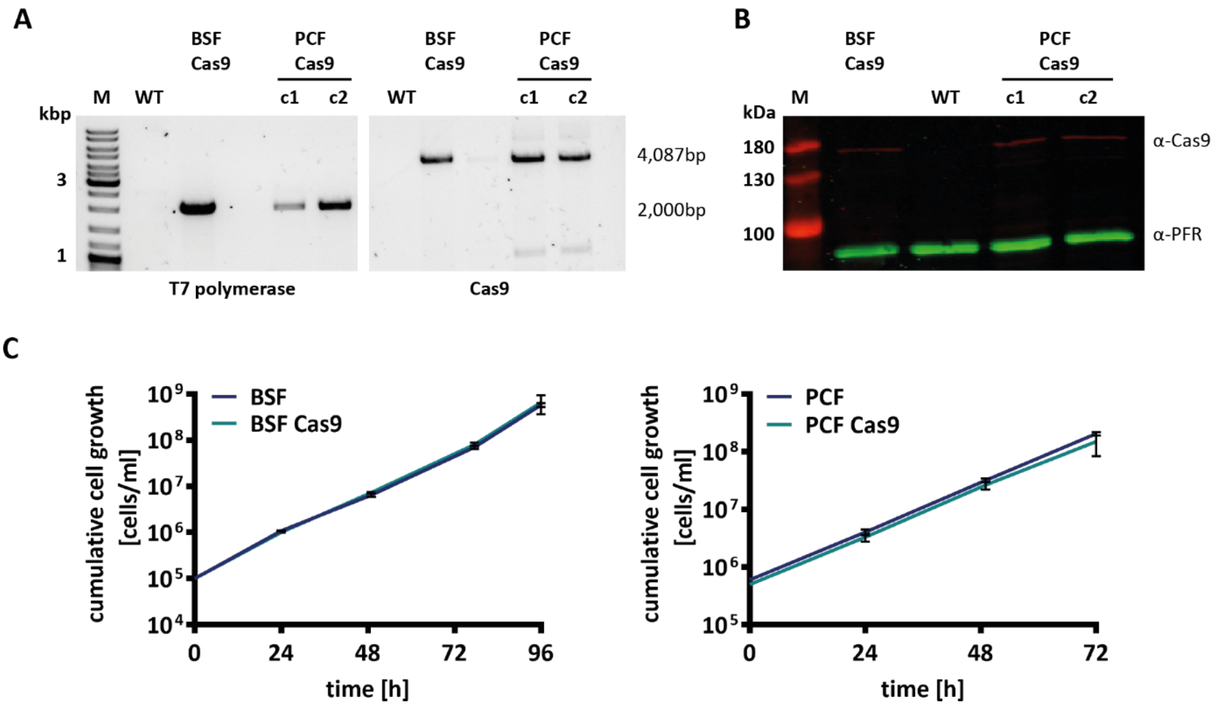


**Supplementary Figure S2: Generation of Cas9-expressing cell lines. (A)** Integration PCR using primers binding within the T7 polymerase and Cas9 genes to confirm construct integration. The genomic DNA of one clone (c1) of BSF T7+Cas9 and two clones (c1 and c2) of PCF T7+Cas9 cells were tested. Wild-type (WT) cell DNA served as a control. **(B)** Western blot showing Cas9 expression in modified BSF and PCF cells. WT served as the control cell line, and PFR was used as the loading control. **(C)** Growth curves of BSF and PCF cells showing that Cas9 expression had no effect on growth (n=3). WT cells (MiTat1.2 wt and 427 strains) served as controls. Error bars are based on the standard deviation (SD).

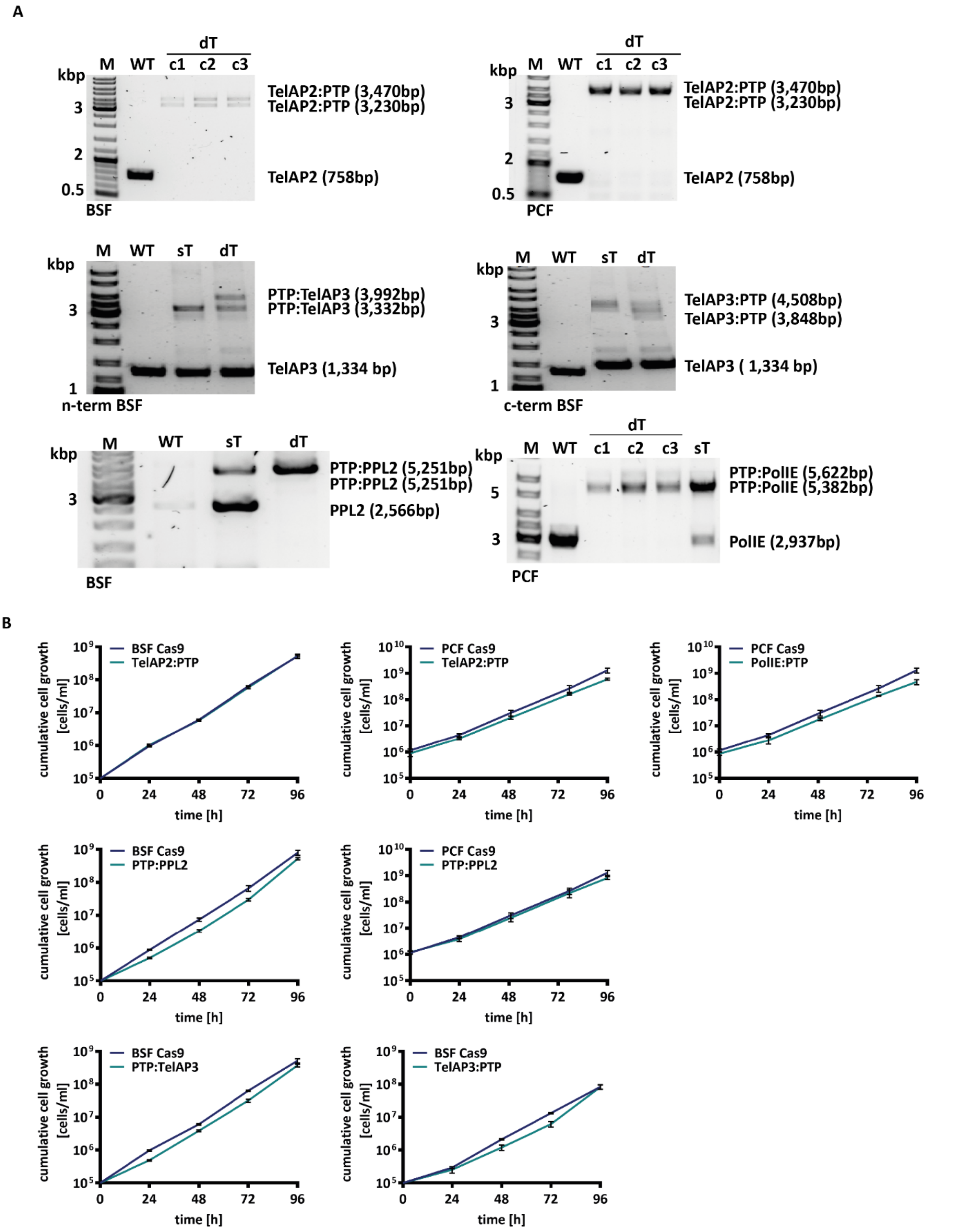


**Supplementary Figure S3: Generation of cell lines for immunoprecipitation. (A)** Validation of tag integration by PCR using primers binding to the 5' and 3'UTRs of the genes of interest. The genomic DNA of different clones (c1-c3), single-tagged (sT) and double-tagged cells (dT) in BSF and PCF cells was used. Wild-type (WT) cell DNA served as a control. **(B)** Growth curves of the tagged cells showing no growth phenotype after PTP tagging. Wild-type cells (BSF Cas9 and PCF Cas9) served as control cell lines.

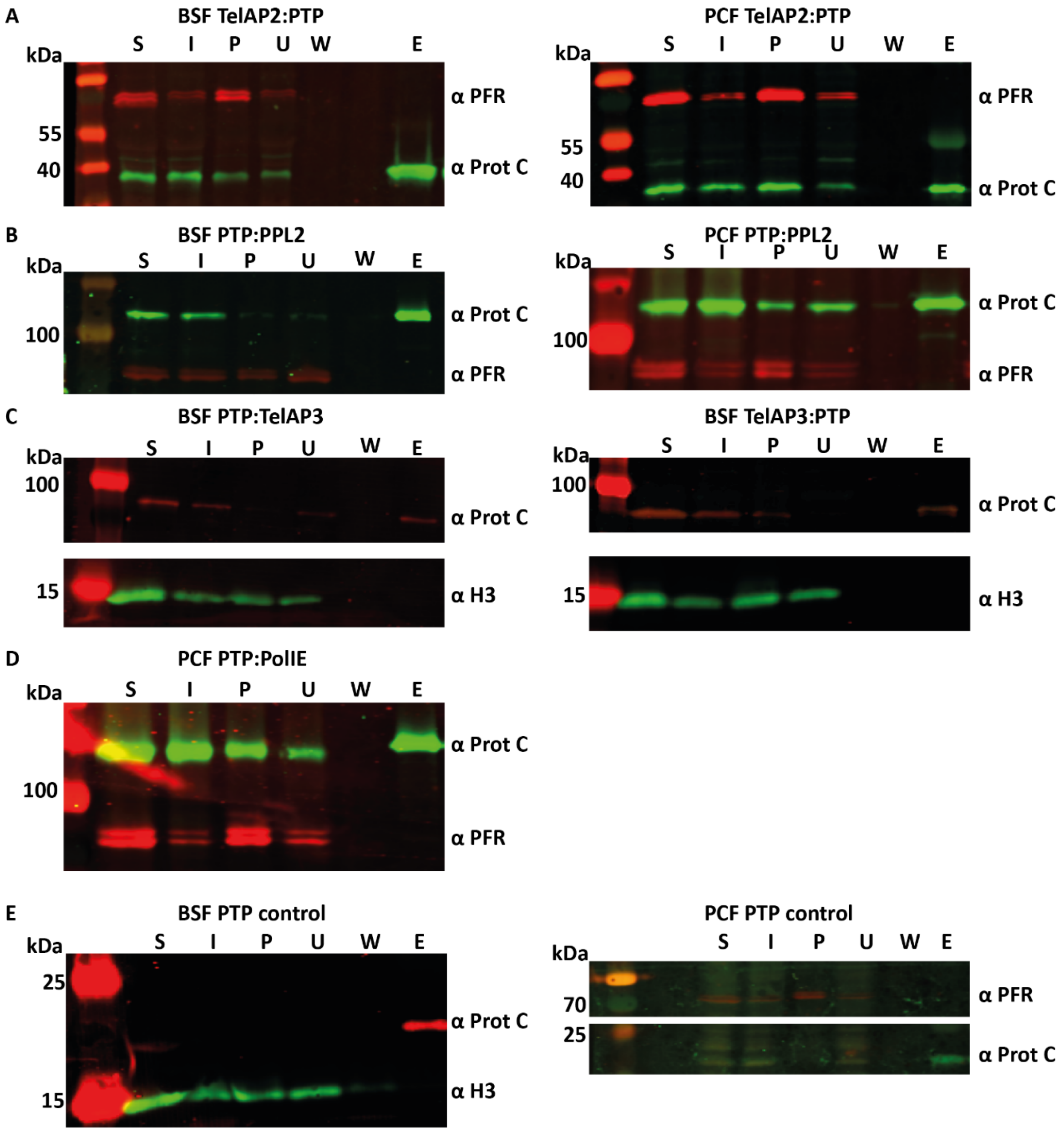


**Supplementary Figure S4: Representative Western blots of the immunoprecipitates. (A)** Representative Western blots of TelAP2:PTP IPs from BSF and PCF cells. Western blots were incubated with anti-protein C and anti-PFR antibodies. The PFR signal served as a loading control. **(B)** Representative Western blots of the IP of PTP:PPL2 in BSF and PCF cells. Western blots were incubated with anti-protein C and anti-PFR antibodies. The PFR signal served as a loading control. **(C)** Representative Western blots of the IP of PTP:TelAP3 and TelAP3:PTP. Western blots were incubated with anti-protein C and anti-H3 antibodies. The H3 signal served as a loading control. **(D)** Representative Western blot of the IP of PTP:PolIE in PCF cells. The Western blot was incubated with anti-protein C and anti-PFR antibodies. The PFR signal served as a loading control. **(E)** Representative Western blots of the immunoprecipitates of ectopically expressed PTPs in BSF and PCF cells. Western blots were incubated with anti-protein C and either anti-PFR or anti-H3 antibodies. The PFR and H3 signals served as loading controls. (S) Start material: whole cell lysate, (I) input material, (P) pellet, (U) unbound material, (W) wash, (E) eluate. A total of 7-8 times more eluate was loaded.

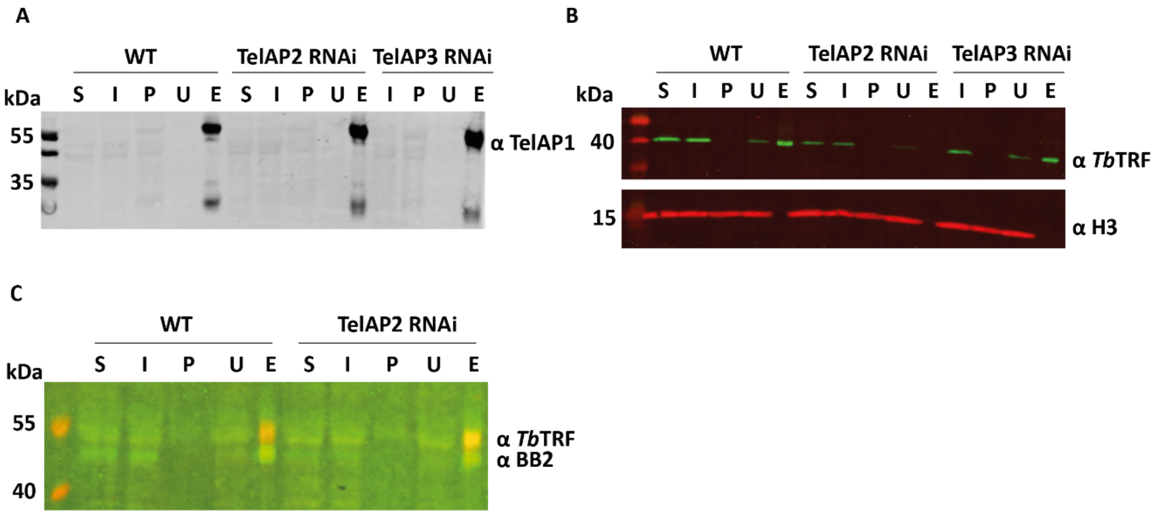


**Supplementary Figure S5: Western blots of immunoprecipitates under RNAi conditions.** IPs with the TelAP1 monoclonal mouse antibody were performed for the WT, TelAP2 RNAi and TelAP3 RNAi cell lines (induced by tetracycline for 48 h prior to IP). **(A)** Representative Western blot of TelAP1 immunoprecipitates incubated with an anti-TelAP1 mouse antibody. **(B)** Representative Western blot of TelAP1 immunoprecipitates incubated with anti-*Tb*TRF and anti-H3 (loading control) antibodies. **(C)** Representative Western blot of *Tb*TRF immunoprecipitates incubated with anti-*Tb*TRF and anti-BB2 antibodies. (S) Start material; whole cell lysate, (I) input material, (P) pellet, (U) unbound fraction and (E) eluate. A total of 7 times more eluate was loaded.

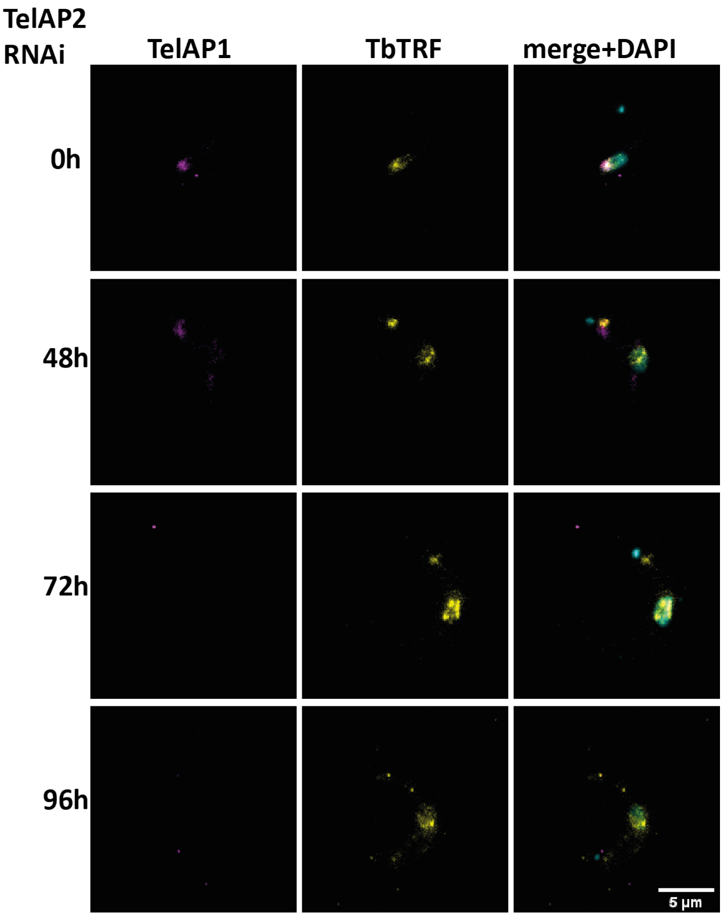


**Supplementary Figure S6: Immunofluorescence of TelAP1 and TbTRF upon TelAP2 knockdown.** Indirect immunofluorescence analysis of uninduced BSF cells and cells 48 h, 72 h and 96 h after TelAP2 depletion via RNAi using monoclonal antibodies against TelAP1 (magenta) and TbTRF (yellow). DNA was stained with DAPI (cyan). Scale bar: 5 µm.

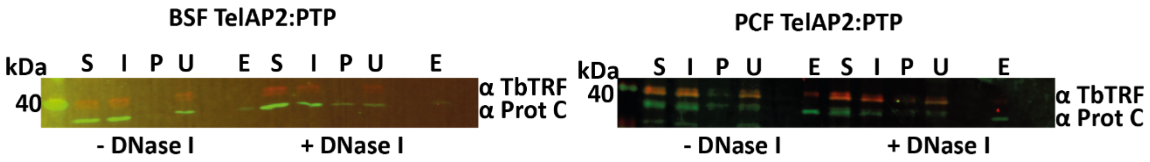


**Supplementary Figure S7: DNA dependence of TelAP2 interactions.** Representative Western blot of BSF and PCF from the IP experiments with and without DNase I treatment. Western blots were incubated with anti-TbTRF and anti-protein C antibodies. (S) Start material: whole cell lysate, (I) input material, (P) pellet, (U) unbound fraction, (W) wash, (E) eluate. A total of 7-fold more eluate material was loaded

1. Aslett M, Aurrecoechea C, Berriman M, Brestelli J, Brunk BP, Carrington M, et al. TriTrypDB: a functional genomic resource for the Trypanosomatidae. Nucleic Acids Res. 2010;38(Database issue):D457-62.

2. Goos C, Dejung M, Janzen CJ, Butter F, Kramer S. The nuclear proteome of Trypanosoma brucei. PLoS One. 2017;12(7):e0181884.
